# Cooption of antagonistic RNA-binding proteins establishes cell hierarchy in *Drosophila* neuro-developmental tumors

**DOI:** 10.1101/353508

**Authors:** Sara Genovese, Raphaël Clément, Cassandra Gaultier, Florence Besse, Karine Narbonne-Reveau, Fabrice Daian, Sophie Foppolo, Nuno Miguel Luis, Cédric Maurange

## Abstract

The mechanisms that govern the hierarchical organization of tumors are still poorly understood, especially in highly heterogeneous neural cancers. Previously, we had shown that aggressive neural tumors can be induced upon dedifferentiation of susceptible intermediate progenitors produced during early development (Narbonne-Reveau et al., 2016). Using clonal analysis, stochastic modelling and single-cell transcriptomics, we now find that such tumors rapidly become heterogeneous, containing progenitors with different proliferative potentials. We demonstrate that tumor heterogeneity emerges from the deregulated transition between two antagonistic RNA-binding proteins, Imp and Syncrip, that switch neural progenitors from a default self-renewing to a differentiation-prone state during development. Consequently, aberrant maintenance of Imp confers a cancer stem cell-like identity as Imp^+^ progenitors sustain tumor growth while being able to continuously generate Syncrip^+^ progenitors. The latter exhibit limited self-renewal likely due to Syncrip-mediated metabolic exhaustion. This study provides an example of how a subverted developmental transition establishes a hierarchical tumor.

## Introduction

The Cancer Stem Cell (CSC)/Hierarchical tumor model postulates that the unlimited self-renewing potential of tumor cells is not a property shared by all cell types within the tumor. Instead, only a subpopulation of the tumor cells, the CSCs, is endowed with the ability to undergo unlimited self-renewal and is therefore responsible for the propagation of tumor growth (Nassar and Blanpain, 2016; Nguyen et al., 2012; Valent et al., 2012). As well as being capable of endless self-renewal, CSCs also generate cellular heterogeneity by producing transit amplifying progenitors (TAPs) that are committed to short term self-renewal and differentiation. The molecular principles that establish the tumor hierarchy and govern the balance between self-renewal or differentiation to generate heterogeneity while maintaining the CSC pool are still poorly defined. Recently, single-cell RNA-seq approaches in various brain tumors have allowed the reconstruction of differentiation trajectories reminiscent to those observed during development, suggesting that neuro-developmental programs may, at least partially, be recapitulated in brain tumors to establish its hierarchy. (Lan et al., 2017; Tirosh et al., 2016; Venteicher et al., 2017). Consequently, it has been proposed that neural cancers may be “locked” into to a developmental program (Azzarelli et al., 2018). This attractive hypothesis could be particularly relevant for pediatric cancers that often originate from growing tissues and require much less mutations than adult cancers to be initiated and become aggressive (Grobner et al., 2018; Scotting et al., 2005; Vogelstein et al., 2013). Therefore, understanding development will help unravelling how its subversion may turn a growing organ into a tumor.

However, unlike developmental programs that should be deployed unidirectionally, it is believed that some tumors exhibit considerable plasticity, with TAPs or more differentiated progeny being able to reverse to a CSC state (Gupta et al., 2011; Meacham and Morrison, 2013). Yet, evidence of rigid vs plastic hierarchy is sparse in neural tumors and the mechanisms implementing uni-or bi-directionality are elusive.

In the face of the complexity of brain development and neural tumors in mammals, simple animal models can represent a powerful alternative to investigate basic and evolutionary conserved principles. The development of the central nervous system (CNS) is undoubtedly best understood in *Drosophila*. The CNS arises from a small pool of asymmetrically-dividing neural stem cells (called neuroblasts (NBs)). NBs divide all along development but terminate before adulthood during metamorphosis (Homem and Knoblich, 2012; Maurange and Gould, 2005). During early larval stages, NBs express the ZBTB transcription factor Chinmo and the RNA-binding protein (RBP) Imp (Narbonne-Reveau et al., 2016; Syed et al., 2017). As NB divide, they sequentially express a series of transcription factors (known as temporal transcription factors) that provides an intrinsic timing mechanism scheduling the systematic silencing of Chinmo and Imp towards mid-larval stages (late L2/early L3)(Maurange et al., 2008; Narbonne-Reveau et al., 2016). Late temporal transcription factors such as Seven-up appear to silence Imp by activating the expression of Syncrip (Syp), another RBP, in early L3 NBs (Ren et al., 2017). By directly or indirectly silencing *Imp*, Syp limits NB self-renewal to ensure that they all undergo terminal differentiation before adulthood. Consequently, loss of Syp leads to NBs that persist in adults and continue dividing (Yang et al., 2017). Together, these studies revealed that the self-renewing potential of NBs during development is limited by the systematic Imp-to-Syp transition scheduled by temporal transcription factors around mid-larval stages.

Aberrant NB amplification can be initiated by perturbing the mechanisms controlling NB asymmetric divisions. For example, upon inactivation of the transcription factor*prospero* (*pros*), NB daughter cells, called Ganglion Mother Cells (GMCs), fail to properly differentiate and reverse to a NB-like state (Bello et al., 2006; Caussinus and Gonzalez, 2005; Knoblich, 2010). We have previously demonstrated that when this event is induced from Chinmo^+^Imp^+^ GMCs that are produced during early larval stages, this leads to aggressive NB tumors (Narbonne-Reveau et al., 2016). Such tumors, that possess an early developmental origin, are therefore a good model to investigate conserved principles underlying pediatric cancers. In addition, we could show that Imp and Chinmo are aberrantly maintained in tumors and are required to propagate their unlimited growth. Interestingly, Imp and Chinmo are only expressed in a subset of the NB-like cells that compose the tumor (Narbonne-Reveau et al., 2016). This has led us to propose that Chinmo^+^Imp^+^ tumor NBs (tNBs) may be equivalent to CSCs, although the heterogeneity within NB tumors needed to be characterized and the existence of a cellular hierarchy remained to be demonstrated.

Here we investigate in details the molecular and cellular heterogeneity of NB tumors. Using *in vivo* lineage analysis combined with numerical modeling, we demonstrate that NB tumors exhibit a rigid hierarchy triggered by the non-systematic Imp-to-Syp transition in dividing tNBs. In addition, using single-cell RNA-seq, we show that Syp induces a metabolic switch in tNBs that is likely to stop growth and promote rapid cell-cycle exit. Our work therefore identifies a subverted developmental transition that induces a tumor “being locked into a developmental program”.

## Results

### Tumors are composed of NBs with either early Imp^+^ or late Syp^+^ identities

Tumors were induced by knocking down *pros* from early larval stages in six NBs located in the ventral nerve cord (VNC) using the *pox*^*n*^-*GAL4*, *UAS-pros*^*RNAi*^ *UAS-GFP*, *UAS-dicer2* system - thereafter referred to as *pox*^*n*^>*pros*^*RNAi*^ (Narbonne-Reveau et al., 2016). *pox*^*n*^>*pros*^*RNAi*^ tumors can persist and expand in adults. We had previously shown that tumor growth in adults requires the co-expression of *Imp* and *chinmo* in a subset of tNBs. Given that a large number of tNBs do not express Chinmo and Imp, we tested for Syp expression in tumors. Immuno-stainings performed a few hours after *pros* knock down (late L2 stage) indicate that tumors develop from a small pool of Chinmo^+^ (Imp^+^) tNBs (Figure 1A).

**Figure 1.**
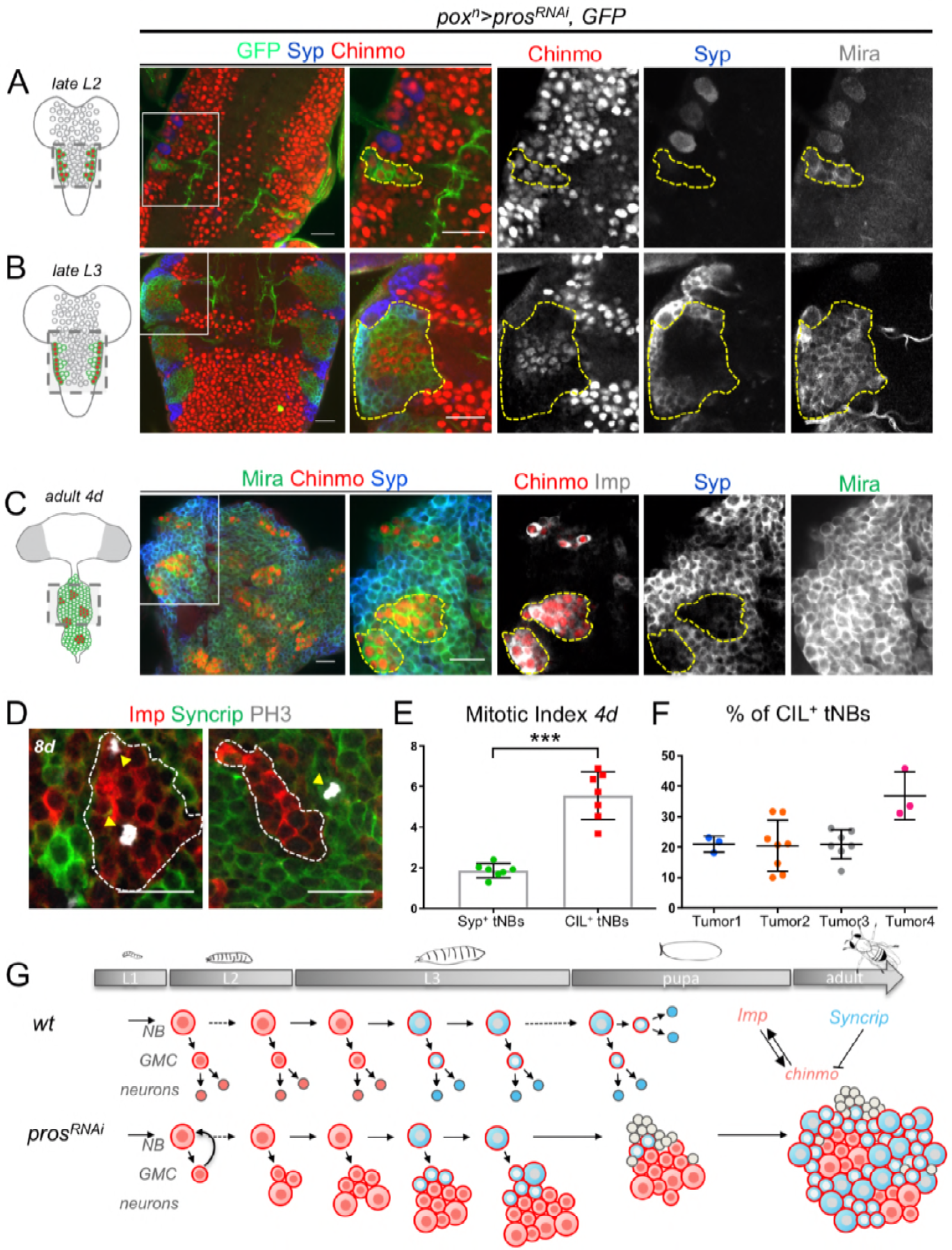
Tumors are composed of two distinct populations of tNBs. Cartoons represent a ventral view of the larval and adult *Drosophila* CNS. Tumor NBs (tNBs) are represented as green circles. Red NBs express *chinmo*. Tumors were induced in L1 by knocking down *pros* in six NBs located in the VNC using the *pox*^*n*^-*GAL4*, *UAS-pros*^*RNAi*^ *UAS-GFP*, *UAS-dicer2* system (*pox*^*n*^>*pros*^*RNAi*^). (**A**) *pox*^*n*^>*pros*^*RNAi*^ tumors are composed exclusively by CIL^+^ tNBs in early larvae (L2). (**B**) In late larvae (L3), *pox*^*n*^>*pros*^*RNAi*^ tumors are composed by two distinct populations of CIL^+^tNBs and Syp^+^tNBs. tNBs are marked with Mira and GFP. Tumors are delimited by dashed lines. (**C**) In adults *pox*^*n*^>*pros*^*RNAi*^ tumors, CIL^+^ tNBs are organized in small clusters surrounded by large fields of Syp^+^ tNBs. tNBs are marked with Mira. (**D**) Mitotic CIL^+^tNBs and Syp^+^tNBs are marked with PH3 in 4-day-old *pox*^*n*^>*pros*^*RNAi*^ tumors. (**E**) Quantification of the mitotic index of CIL^+^ tNBs (n=7 VNC) and Syp^+^ tNBs (n=7 VNC) in 4-day-old *pox*^*n*^>*pros*^*RNAi*^ tumors. *P* = 0.0006. CIL^+^ tNBs have a higher mitotic index than Syp^+^ tNBs in adult *pox*^*n*^>*pros*^*RNAi*^ tumors. (**F**) Proportion of CIL^+^tNBs in *pox*^*n*^>*pros*^*RNAi*^ tumors from 2-5 day-old flies (n=4 VNC, plotted individually; data points are planar sections at increasing depths from the tumor surface). (**G**) Scheme depicting tumor initiation and progression upon *pros* knock down from early larval development. Scale bars, 20 μm.

No Syp^+^ tNBs are present at this stage. However, from mid to late L3, a distinct population of Syp^+^ tNBs emerges within tumors (Figure 1B). Thus, Syp^+^ tNBs appear in tumors at about the time when normal NBs undergo the Imp→Syp switch. In adults, Chinmo^+^Imp^+^ tNBs are encountered in small clusters surrounded by large fields of Syp^+^ tNBs (Figure 1C). Thus, the initial pool of Chinmo^+^Imp^+^ tNBs evolves to generate two distinct compartments within tumors that are reminiscent of early and late stage NBs during development. Interestingly, the complementary Imp and Syp stainings in adult tumors appear to encompass all tNBs. Both the Imp^+^ and Syp^+^ compartments show mitotic activity (Figure 1D), although the mitotic index is significantly lower in the Syp^+^ compartment (Figure 1E). Then, we used Tissue Analyzer, a segmentation tool, to precisely quantify the proportion of Imp^+^ and Syp^+^ tNBs within tumors (Aigouy et al., 2016). While the number of tNBs continuously increases in adults over time (for example, the volume of tNBs increase by more than 10 fold between 1-day and 6-day-old adults) (Narbonne-Reveau et al., 2016), we found that the proportion of Imp^+^ tNBs remained relatively stable, between 20 to 30% of all tNBs in different tumors from 2-to-5-day-old adults (Figure 1F). Thus, Imp^+^ tNBs exhibit a higher mitotic index but they rapidly become a minority compared to Syp^+^ tNBs (Figure1F,G). This cannot be explained by preferential apoptosis or neuronal differentiation of Imp^+^ tNBs as these events are rare and usually more frequent among the Syp+ tNBs (Figure 1-figure supplement 1A-C). In conclusion, tumors rapidly evolve from a homogenous population of Imp^+^ tNBs in early larvae to an heterogeneous population of Imp+ and Syp+ tNBs with a ≈20/80% ratio in adults.

### NB tumors follow a rigid hierarchical scheme

To investigate in details the rules governing the population dynamic of Imp^+^ and Syp^+^ tNBs, we designed lineage analysis experiments *in vivo*. We used the Flybow technique combined with our *pox*^*n*^>*pros*^*RNAi*^ system to generate random RFP-labeled clones in tumors (Hadjieconomou et al., 2011). Clones were randomly induced in 2-day-old adults containing*pox*^*n*^>*pros*^*RNAi*^ tumors. All tNBs are labelled either with CD8-GFP or with CD8-Cherry. Individual CD8-Cherry^+^ clones were examined 8 hours, 2 days, 4 days and 8 days after clonal induction (ACI) (Figure 2A). We used Chinmo as a marker for Imp+ tNBs and its absence as a marker for Syp^+^ tNBs. Along this time lapse, we could observe three categories of clones: clones composed of Chinmo^+^Imp^+^ tNBs only (further termed CI clones), MIXED clones composed of both Chinmo^+^Imp^+^ and Syp^+^ tNBs (they therefore contain at least 2 tNBs), and clones composed of Syp^+^ tNBs only (further termed SYP clones) (Figure 2C). The existence of clones only composed of Imp^+^ or Syp^+^ tNBs is consistent with the ability of both types of tNBs to undergo self-renewing divisions. In addition, the existence of MIXED clones shows that Imp^+^ and Syp^+^ tNBs may derive from a common precursor of either identity. This clonal relationship could be indicative of a hierarchical scheme. In addition, clones display a large heterogeneity in their size at 8 days ACI (Figure 2A,B), implying that all tumor cells do not possess the same proliferation potential, a characteristic of hierarchical tumors (Nassar and Blanpain, 2016). Interestingly, we observed that the proportion of the different categories of clones is dynamic over the period of 8 days. The proportion of CI clones exhibits a rapid decrease that is paralleled with an increase in the proportion of MIXED clones and a slight increase in the proportion of SYP clones (Figure 2D). We sought to investigate whether this dynamics could reflect a rigid or plastic hierarchy within the tumor.

**Figure 2.**
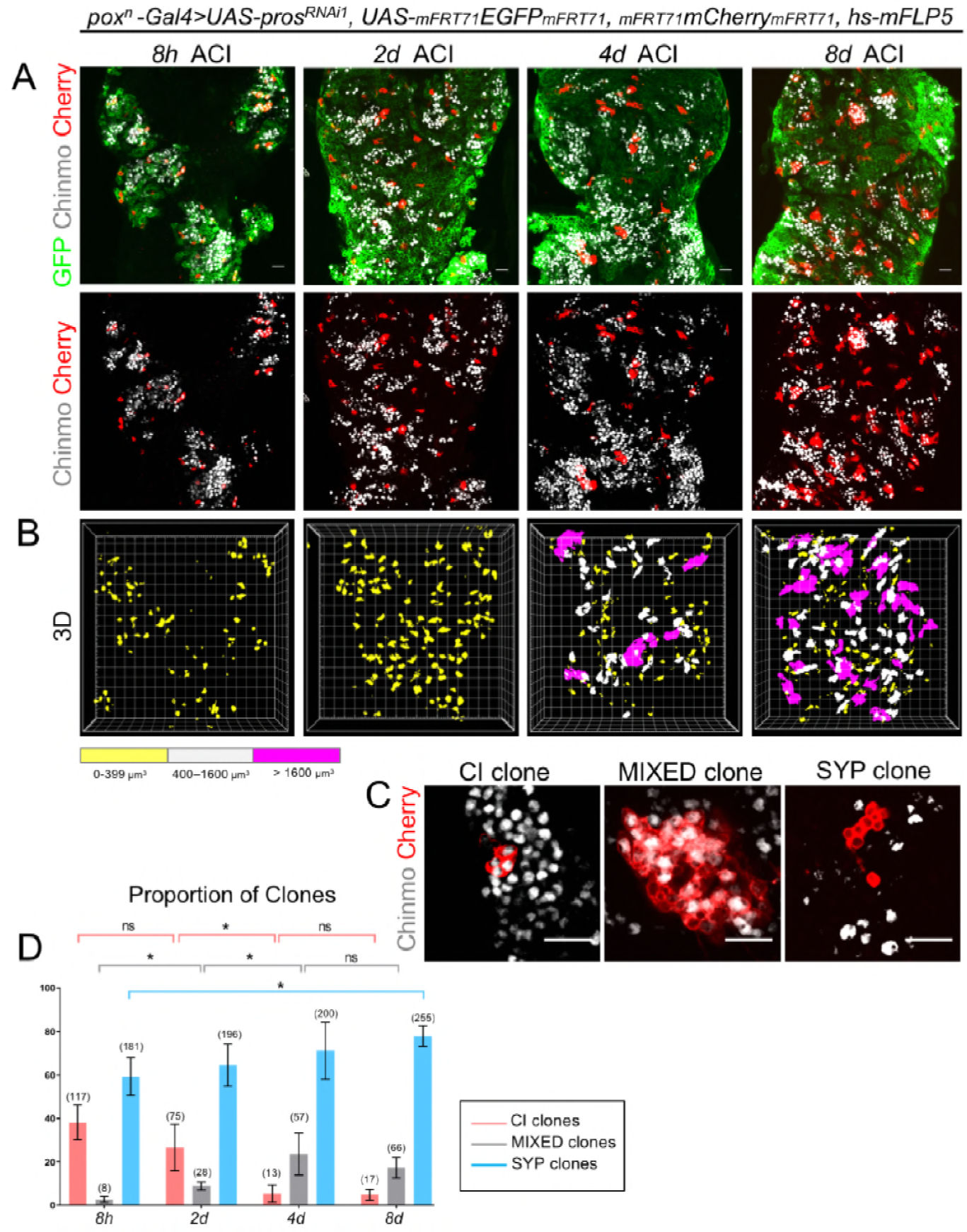
Clonal analysis in tumors. (**A**) Clones are labelled with Cherry and observed 8 hours (8h), 2 days (2d), 4 days (4d) and 8 days (8d) after clonal induction (ACI) in *pox*^*n*^ >*pros*^*RNAi*^ tumors. Cherry^−^ tNBs are GFP^+^. Images represent one confocal section. (**B**) 3D projections of clones 8h, 2d, 4d and 8d ACI. The color-code labels clones according to their volume. (**C**) Three categories of clones can be identified in *pox*^*n*^>*pros*^*RNAi*^ tumors. Chinmo^+^Imp^+^ tNBs are identified by the presence of Chinmo. Syp^+^ tNBs are identified by the absence of Chinmo. (**D**) Proportion of CI (red), MIXED (grey) and SYP (blue) clones 8h, 2d, 4d and 8d ACI. Proportion of CI clones 8h ACI (n=117 clones form 4 VNCs); 2d ACI (n=75 clones from 4 VNCs); 4d ACI (n=13 clones from 5 VNCs); 8d ACI (n=17 clones from 5 VNCs). *P* between CI clones at 8h and 2d ACI -> *P*_Ci8H/2d_ = 0.2; *P*_CI2d/4d_ = 0.016; *P*_CI4d/8d_ = 0.88. Proportion of MIXED clones 8h ACI (n=8 clones from 4 VNCs); 2d ACI (n=28 clones from 4 VNCs); 4d ACI (n=57 clones from 5 VNCs); 8d ACI (n=66 clones from 5 VNCs). *P*_MIXED 8h/2d_ = 0.029; *P*_MIXED2d/4d_ = 0.016; *P*_MixED4d/8d_ = 0.31. Proportion of SYP clones 8h ACI (n=181 clones from 4 VNCs); 2d ACI (n= 196 clones from 4 VNCs); 4d ACI (n=200 clones from 5 VNCs); 8d ACI (n=255 clones from 5 VNCs). *P*_Syp8H/8d_ = 0.016. Scale bars, 20 μm.

For this purpose, we designed a stochastic numerical model of clone growth (see Material and Methods for details). The model follows a simple algorithm (Figure 3A). Each clone starts from a single tNB. In the model, Chinmo^+^Imp^+^ tNBs are referred to as C cells while Syp^+^ tNBs are referred to as S cells. The initial tNB can either be Chinmo^+^Imp^+^ (probability *p*_*c*_) or Syp^+^ (probability 1 − *p*_*c*_). At each numerical time step, each cell in the clone (initially one) has a given probability to divide, set by its division time. Upon division, Chinmo^+^Imp^+^ tNBs can either duplicate (C→CC), generate two Syp^+^ tNBs (C→SS), or undergo asymmetric division (C→CS). Similarly, Syp^+^ tNBs can either duplicate (S→SS), generate two Chinmo^+^Imp^+^ tNBs (S→CC), or undergo asymmetric division (S→CS). Each new tNB has a probability to be quiescent. In our *pox*^*n*^ >*pros*^*RNAi*^; *Flybow* combination, the Cherry is not maintained in differentiated neuronal progeny possibly generated within clones. Therefore, to better fit to our clonal analysis, we did not consider cell death or differentiation into neurons in the model. In modeling terms, we asked whether hierarchical or plastic schemes of cell divisions are compatible with the experimental observations. We first tested three simplified scenarios, in which we neglected quiescence and asymmetric divisions, and assumed that C and S cells have the same division time. In each case, we simulated the growth of 1000 model clones and plotted the proportion of clones in each category. In the first scenario (high plasticity), Chinmo^+^Imp^+^ tNBs have 50% chance to duplicate (symmetric self-renewing divisions C→CC), and 50% chance to divide into two Syp^+^ tNBs (C→SS). Similarly, Syp^+^ tNBs have 50% chance to duplicate (S→SS), and 50% chance to divide into two Chinmo^+^Imp^+^ tNBs (S→CC). Unlike our experimental observations, this non-hierarchical scenario leads to a symmetric outcome, in which all clones rapidly become mixed, while the proportion of CI and SYP clones rapidly decays to zero (Figure 3B, left panel). In the second scenario (rigid hierarchy), Chinmo^+^Imp^+^ tNBs still have 50% chance to duplicate (C→CC), and 50% chance to divide into two Syp^+^ tNBs (C→SS) but Syp^+^ tNBs can only self-renew (S→SS). This hierarchical scheme leads to a fully different outcome, with a large proportion of clones remaining only composed of Syp^+^ tNBs (SYP clones), while the proportion of clones only composed of Chinmo^+^Imp^+^ tNBs (CI clones) rapidly decays, which is consistent with our experimental observations (Figure 3B, middle panel). In the third scenario (low plasticity), we maintain this hierarchy, but also give Syp^+^ tNBs a small probability (1%) to generate Chinmo^+^Imp^+^ tNBs (S→CC). This is sufficient to drastically change the outcome of the simulations, as it prevents the maintenance of SYP clones (Figure 3B, right panel). These three model scenarios combined to our experimental observations argue for a rigid hierarchy between tNBs, with a negligible or zero probability for Syp^+^ tNBs to generate Chinmo^+^Imp^+^ tNBs, as such divisions prevent the long-term maintenance of SYP clones observed *in vivo*.

**Figure 3.**
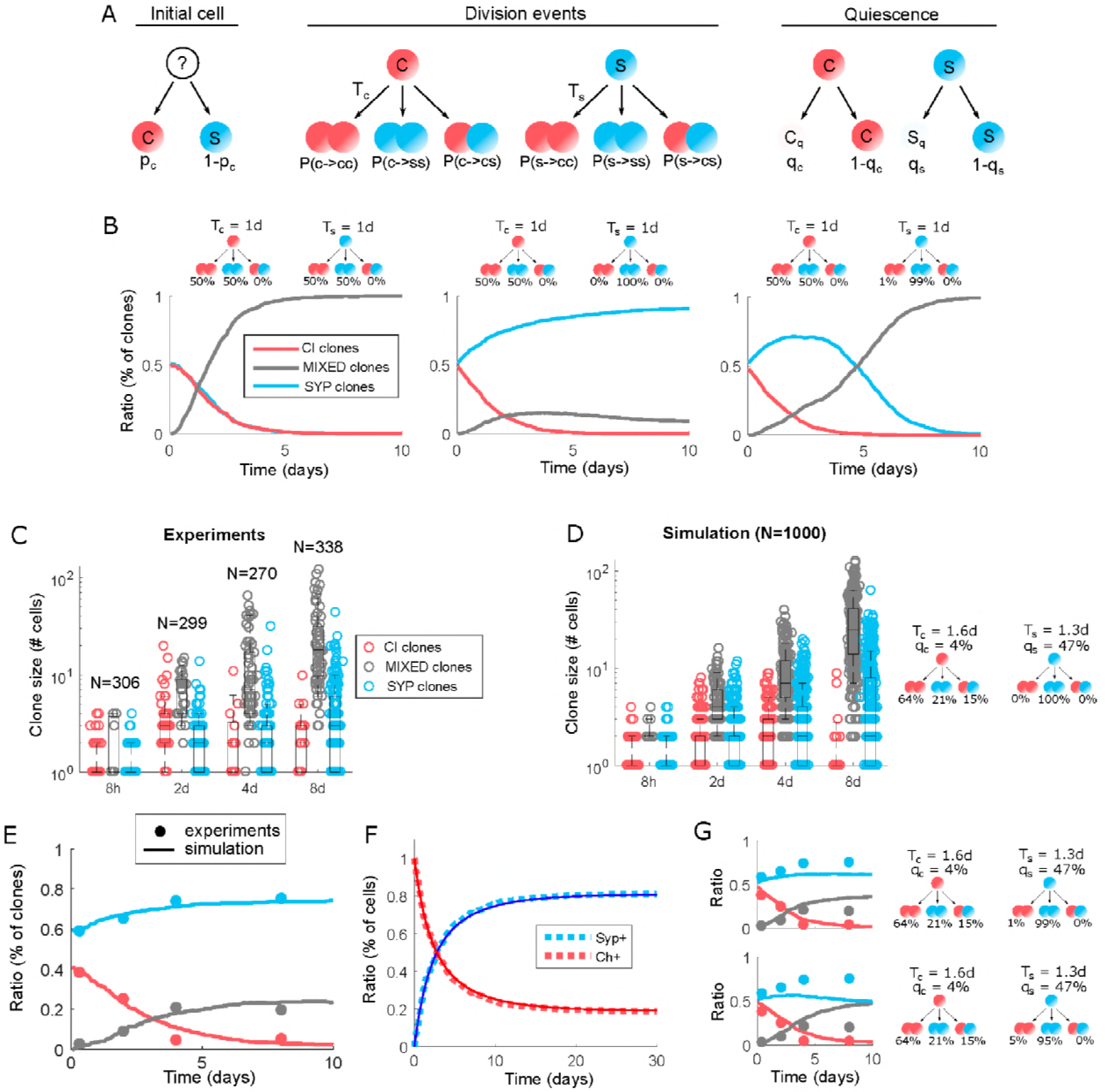
Stochastic model of clone growth. (**A**) Cartoon of the stochastic clone model. (**B**) Proportion of clones in each category in three extreme model scenarios: high plasticity (left panel), rigid hierarchy (middle panel), or low plasticity = loose hierarchy (right panel). Cartoons above graphs represent the division probabilities used to generate each graph. (**C**) Clone sizes for each category calculated from the experiments (**D**) Clone sizes for each category using the set of parameters minimizing the error. Center lines of boxes show the medians, boxes limits indicate the 25th and 75th percentiles, and whiskers limits indicate the 10th and 90th percentiles. (**E**) Proportion of clones overtime in each category using the set of parameters minimizing the error. Solid lines represent the simulations (n=1000 clones), dots represent experimental measurements. (**F**) Proportion of Syp^+^ (blue) vs Chinmo^+^Imp^+^ (red) tNBs over time using the set of parameter values determined by the error minimization. Dashed lines represent a stochastic simulation (clone model), thin lines represent the prediction of the deterministic model. (**G**) Proportion of clones in each category using the set of parameters minimizing the error while allowing a small chance of reverse division from S to C (top: 1%, bottom: 5%).

### Quantitative analysis of hierarchical tumor growth

We then sought to use the rigid hierarchical model as a basis to investigate more quantitatively the parameters that control tumor growth. Interestingly, significant differences in average clone size can be observed between the three categories at the different time points (Figure 3C). SYP clones grow slower than MIXED clones, suggesting either that Syp^+^ tNBs have a slower cell-cycle speed or a self-renewing potential limited by a high probability to exit the cell-cycle - similar to Syp^+^ NBs during development. CI clones tend to completely stop growing, suggesting that the few clones that remain within this category 8 days ACI are composed by quiescent Chinmo^+^Imp^+^ tNBs. We now take into account non-zero probabilities *q_s_* and *q_c_* for new S cells and C cells to be quiescent, a non-zero probability for C cells to undergo asymmetric divisions, and distinct division times for C and S cells. Thanks to a series of measurements, we could reduce the number of independent parameters to 2 (see methods): the probability *P*(*c* → *cc*), and the division time *T*_*s*_ for S cells. We used these as free parameters to fit the model to the measurements of clone sizes and compositions at 8h, 2, 4 and 8 days ACI. To that end, we computed error maps for clone composition and clone size in the (*Pc* → *cc*, *T*_*s*_) plane, and eventually a combined error map (Figure 3-figure supplement 1). We find that the error is minimized for *P*(*c* → *cc*)) = 0.64 and *T*_*s*_ = 1.3 days, from which we determine all the other parameters. Notably, Syp^+^ tNBs have a much higher probability (47%) to enter quiescence than Chinmo^+^Imp^+^ tNBs (4%), consistent with the slower growth of SYP clones.

Using the set of parameters determined by the fit, we found an excellent agreement between experiments and simulations, for both clone sizes (Figure 3C,D) and clone compositions (Figure 3E). Starting from a homogenous pool of Chimo^+^Imp^+^ tNBs, we also find that the overall proportion of Syp^+^ tNBs plateaus to about 80% of the tumor (Figure 3F). Note however that the final proportion of the two types of tNBs is largely affected by the probability for Syp^+^ tNBs to enter quiescence (Figure 3-figure supplement 2). These simulations are consistent with our observation that tumors evolve from homogeneity to stable heterogeneity (Figure 1).

With these parameters, allowing a small probability for S→CC reverse divisions significantly affects the simulations outcome, leading to an important loss of SYP clones and a concomitant increase of MIXED clones (Figure 3G). This further demonstrates that reverse divisions from Syp^+^ tNBs to Chinmo^+^Imp^+^ tNBs are either very low or inexistent.

Altogether, our clonal analysis demonstrates that NB tumors are strongly hierarchical. Chinmo^+^Imp^+^ tNBs are at the top at the hierarchy. They sustain tumor growth through a preference for symmetric self-renewing divisions. They can also undergo asymmetric or symmetric differentiation divisions in order to generate Syp^+^ tNBs, albeit with a lower probability. Syp^+^ tNBs exhibit a high propensity for rapid cell-cycle exit and cannot generate Chinmo^+^Imp^+^ tNBs. Consequently, Syp^+^ tNBs cannot generate large and heterogeneous clones in tumors unlike Chinmo^+^Imp^+^ tNBs. These characteristics confer CSC-like and TAP-like properties to Chinmo^+^Imp^+^ tNBs and Syp^+^ tNBs respectively.

### Loss of Syp abrogates hierarchy

To investigate the function of Syp within the tumor context we knocked it down in *pox*^*n*^>*pros*^*RNAi*^ tumors from their initiation. Interestingly, the subsequent *pros*^*RNAi*^; *Syp*^*RNAi*^ tumor became mostly composed of Chinmo^+^Imp^+^ tNBs (Figure 4A). Thus, Syp knock-down leads to tNBs that fail to silence *chinmo* and *Imp* leading to the dramatic amplification of Chinmo^+^Imp^+^ tNBs (Figure 4B). Our numerical model predicts that abrogation of the Imp→Syp transition in tNBs leads to increased tumor growth rate (Figure 4D). To investigate whether this is also observed *in vivo*, we measured the volume of*pros*^*RNAi*^; *Syp*^*RNAi*^ tumors in adults. In 1-day-old adults, a 3.9-fold increase in tumor volume was observed in *pros*^*RNAi*^; *Syp*^*RNAi*^ compared control *pros*^*RNAi*^ tumors (Figure 4C, E). In addition, over-expressing *Syp* within *pros*^*RNAi*^ tumors blocked tumor growth in adults (Figure 4C, G).

Next, we investigated whether Imp and Chinmo are necessary for the increased tumor growth rate induced by the loss of Syp. Knock-down of both Syp and Imp in *pros*^*RNAi*^ tumors arrests tumor growth in 4-day-old adults (Figure 4C, F). Interestingly, *pros*^*RNAi*^, *Syp*^*RNAi*^, *Imp*^*RNAi*^ tumors lacked or exhibited low levels of Chinmo expression (Figure 4C and Figure 4-figure supplement 1A), consistent with our previous finding that *chinmo* expression relies on Imp (Narbonne-Reveau et al., 2016). Thus, the overgrowth phenotype observed in *pros*^*RNAi*^, *Syp*^*RNAi*^ tumors is mediated by the Chinmo/Imp module. Altogether, our data indicate that tumor heterogeneity is not required for tumor growth. Instead, Syp induces hierarchy by limiting tNB self-renewal, and restricts tumor growth by silencing the Chinmo/Imp module.

### Imp and Syp post-transcriptionally regulate *chinmo* to regulate NB growth and self-renewal

It has recently been shown that Imp and Syp antagonistically regulate *chinmo* in a subset of neurons in the brain, although their mode of action is unclear (Liu et al., 2015). Given that Chinmo is silenced by Syp in tNBs and requires Imp for its expression, we investigated whether chinmo mRNA could be a direct target of both Imp and Syp. As revealed by affinity pull-down assay performed using *in vitro* synthesized *chinmo* 5’ and 3’UTRs, both Imp and Syp can associate with the UTRs of *chinmo* mRNA (Figure 5A). The binding of Syp to the 5’UTR of *chinmo* mRNA is consistent with the previous observation that *chinmo* 5’UTR is important to mediate post-transcriptional silencing (Zhu et al., 2006). This suggests that Imp and Syp may exert their antagonistic activity on *chinmo* expression by binding to the UTRs of its mRNA. In addition, we tested the function of *chinmo* in NBs during development. While the knock down of *Syp* in VNC NBs (using *nab>Gal4*, *UAS-dicer2*, *UAS-GFP*, *UAS-Syp*^*RNAi*^) induced Chinmo and Imp maintenance and persistence of dividing NBs in adults (Figure 5C), we find that the double knock-down of Syp and Imp led to NBs that failed to maintain Chinmo up to late developmental stages (Figure 5E). In addition, NBs were smaller, less numerous and proliferative in adults, often displaying a cytoplasmic extension reminiscent of quiescent NBs (Figure 5E-H) (Chell and Brand, 2010). Similar results are obtained with the double knock-down of *Syp* and *chinmo*, and *Imp* expression was lost (Figure 5D,F,G,H). Thus, Chinmo and Imp form a positive feedback loop that is active during both development and tumorigenesis and is required to propagate long-term NB growth and self-renewal. Activation of Syp, during late developmental stages or in tNBs, exhausts the self-renewing potential of (t)NBs by silencing the Chinmo/Imp module (Figure 5I).

**Figure 4.**
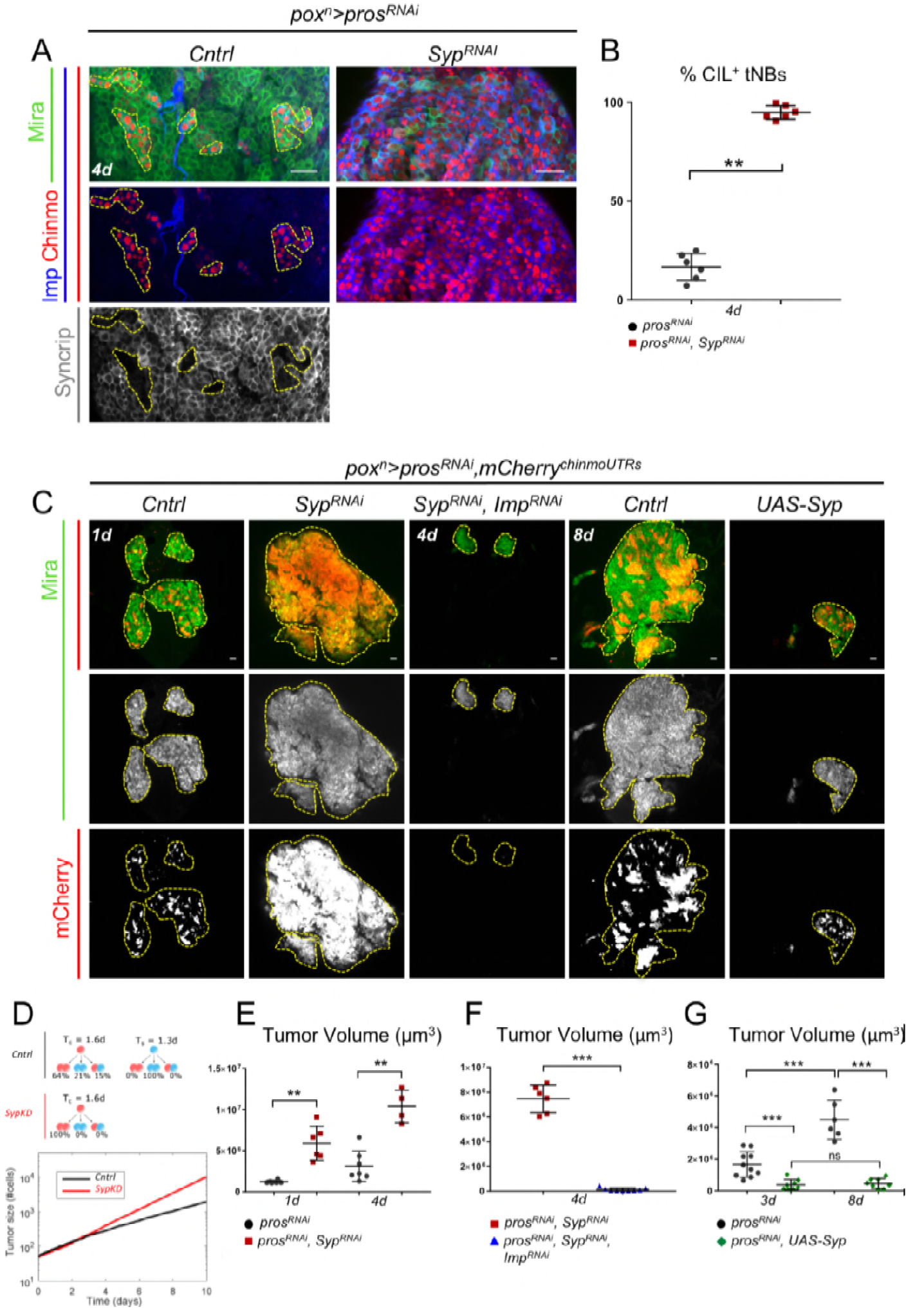
Syp suppresses tumor growth by repressing the Chinmo/Imp modules. Tumors were induced in LI using the *pox*^*n*^ >*pros*^*RNAi*^ system. (**A**) Syp knock-down *in pox*^*n*^>*pros*^*RNAi*^ tumors triggers a dramatic increase of Chinmo^+^Imp^+^tNBs. Clusters of Chinmo^+^Imp^+^tNBs, marked with Chinmo and Imp, are delimited by dashed lines in the Control condition. tNBs are marked by Mira. (**B**) Proportion of Chinmo^+^Imp^+^ tNBs in *pox*^*n*^>*pros*^*RNAi*^ (n=6 VNC) and *pox*^*n*^ >*pros*^*RNAi*^, *Syp*^*RNAi*^ (n=6 VNC) in 4-day-old tumors. *P*= 0.0022. (**C**) Syp knockdown (*Syp*^*RNAi*^) *in pox*^*n*^>*pros*^*RNAi*^ tumors strongly enhances tumor growth in 1 - day-old adults compared to control tumors (*Cntrl*); Adult tumors fail to grow when Imp is knocked down (*Imp*^*RNAi*^); overexpression of *Syp* (*UAS-Syp*) in *pox*^*n*^ >*pros*^*RNAi*^ tumors arrests tumor growth. Tumors are delimited by dashed lines. The *UAS-mCherry*^*chinmoUTRs*^ is expressed in tumors where it is post-transcriptionally regulated reflecting *chinmo* expression (Figure 4-figure supplement 1B). Thus, Chinmo^+^Imp^+^ tNBs are identified by the presence of mCherry. (**D**) Simulation of tumor growth (number of cells) in *pros*^*KD*^ (*Cntrl*-black) and *pros*^*KD*^, *Syp*^*KD*^ (red) tumors. Quiescence probabilities are those set by the error minimization. (**E**) Tumor volume in *pox*^*n*^>*pros*^*RNAi*^ (n=6 VNC) and *pox^*n*^>*pros*^*RNAi*^, *Syp*^*RNAi*^* (n=6 VNC) in 1-day-old adults. *P*=0.002. Tumor volume in *pox*^*n*^>*pros*^*RNAi*^ (n=7 VNC) and *pox*^*n*^>*pros*^*RNAi*^, *Syp*^*RNAi*^ (n=4 VNC) in 4-day-old adults. *P*=0.0061. (**F**) Tumor volume in *pox*^*n*^>*pros*^*RNAi*^, *Syp*^*RNAi*^ (n=6 VNC) and *pox*^*n*^>*pros*^*RNAi*^, *Imp*^*RNAi*^, *Syp*^*RNAi*^(n=9 VNC) in 4-day-old adults. *P*=0.0004. (**G**) Tumor volume in 3-day-old (n=10 VNC) and 8-day-old (n=6 VNC)*pox*^*n*^>*pros*^*RNAi*^ tumors. *P*=0.00025. Tumor volume in *pox*^*n*^>*pros*^*RNAi*^ (n=10 VNC) and *pox*^*n*^>*pros*^*RNAi*^, *UAS-Syp* (n=8 VNC) in 3-day-old adults. P=0.0005. Tumor volume in *pox*^*n*^>*pros*^*RNAi*^ (n=6 VNC) and *pox*^*n*^>*pros*^*RNAi*^, *UAS-Syp* (n=8 VNC) in 8-day-old adults. *P*=0.00067. Tumor volume in 3-day-old (n=8 VNC) and 8-day-old (n=8 VNC) *pox*^*n*^>*pros*^*RNAi*^, *UAS-Syp* tumors. *P*=0.7209. Scale bars, 20 μm.

**Figure 5.**
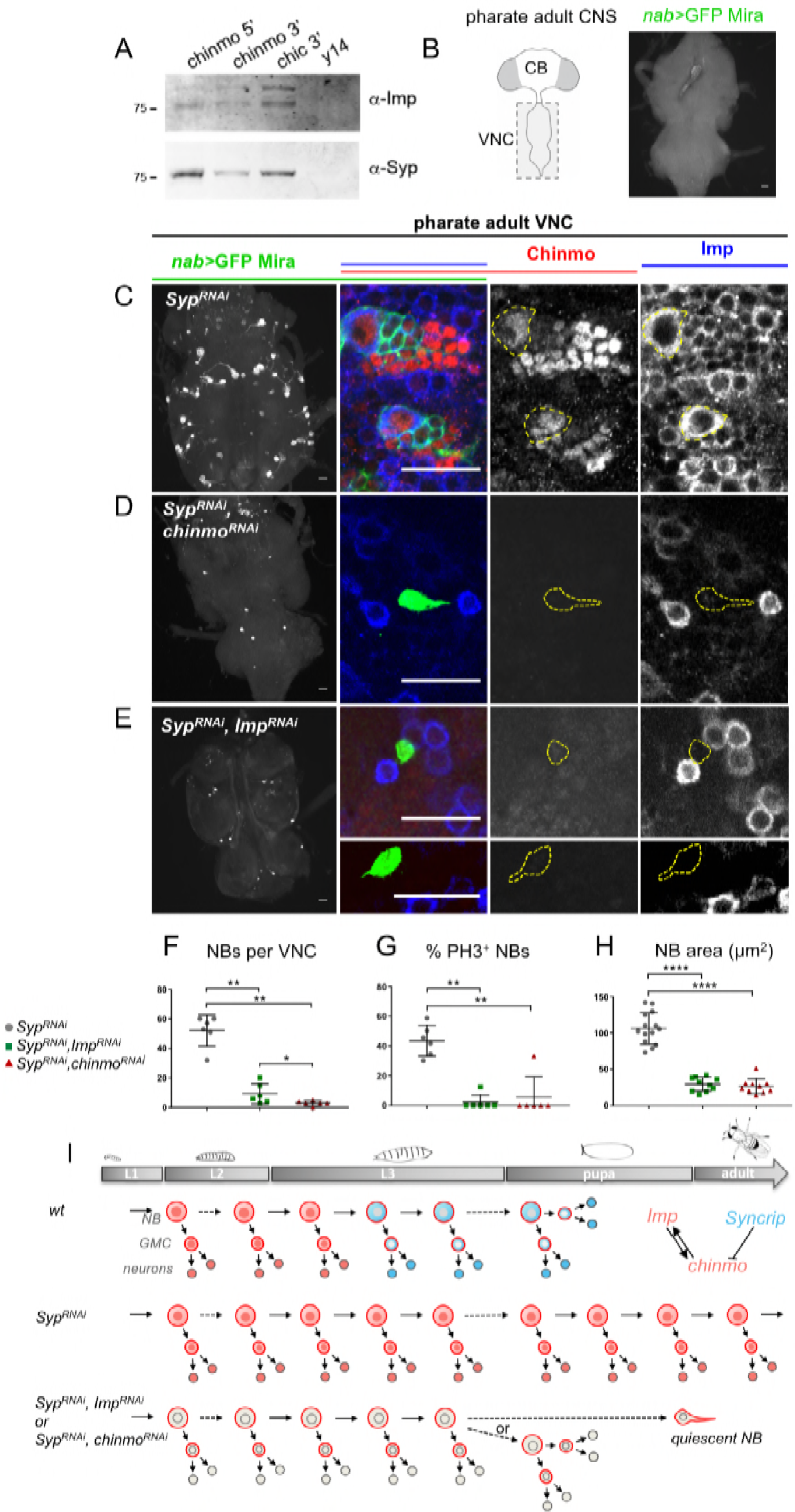
Syp silences Chinmo and Imp to limit NB self-renewal during development. (**a**) RNA affinity pull-down assay using biotinylated RNAs incubated with wild-type adult brain extracts. Proteins recovered in the bound fractions were visualized by western blot analysis. The 3′ UTR of *chickadee* (*chic*) and the coding sequence *of y 14* were used as positive and negative controls respectively, (**b**) Cartoons represent a ventral view of the adult *Drosophila* central nervous system (CNS). The *nab-GAL4* driver is active in all NBs located in the ventral nerve cord (VNC). ln *nab>GaI4*, *UAS-dicer2*, *UAS-GFP* (*nab*) pharate adults, GFP labels NBs and their recent progeny. NBs, marked with GFP and Miranda (Mira) are not present in the VNC of *wt* pharate adults, (**c**) Syp knock-down (*nab>Syp*^*RNAi*^) from early larval stages induces the persistence of active *Chinmo*^+^ and *Imp*^+^ NBs in the VNC of pharate adults, (**d**) Most of the VNC NBs lacking both Syp and chinmo (*nab>Syp*^*RNAi*^, *chinmo*^*RNAi*^) are eliminated before the pharate adult stage. The few persisting NBs exhibit a smaller size and cytoplasmic extensions. They don’t express Chinmo and Imp. (**e**) Most of the VNC NBs lacking both Syp and Imp(*nab>Syp*^*RNAi*^, *Imp*^*RNAi*^) are eliminated before the pharate adult stage. The few persisting NBs exhibit a smaller size and cytoplasmic extensions. They don’t express Chinmo and Imp. (**f**) Quantification of the number of NBs per VNC in *nab>Syp*^*RNAi*^ (n= 6 VNC), nab>*Syp*^*RNAi*^ *Imp*^*RNAi*^ (n= 6 VNC) and *nab>Syp*^*RNAi*^, *chinmo*^*RNAi*^(n= 6 VNC). *P*_*SypKD/SypKD,ImpKD*_= 0.002. *P*_*SypKD/SypKD,chinmoKD*_= 0.002. *P*_*SypKD/SypKD,ImpKD/SypKD/SypKD,chinmoKD*_= 0.039. (**g**) Quantification of proliferating NBs, marked with PH3, in nab>*Syp*^*RNAi*^(n= 6 VNC), nab>*Syp*^*RNAi*^, *Imp*^*RNAi*^ (n= 6 VNC) and nab>*Syp*^*RNAi*^, *chinmo*^*RNAi*^ (n= 6 VNC). *P*_*SypKD/SypKD,ImpKD*_ = 0.002. *P*_*SypKD/SypKD,chinmoKD*_ = 0.004. (**h**) Quantification of the NB area in nab>*Syp*^*RNAi*^(n= 14 NBs), nab>*Syp*^*RNAi*^ *Imp*^*RNAi*^ (n= 10 NBs) and nab>*Syp*^*RNAi*^, *chinmo*^RNAi^ (n= 10 NBs). *P*_*SypKD/SypKD,ImpKD*_ = <0.0001. *P*_*SypKD/SypKD,chinmoKD*_ = <0.0001. (**i**) Scheme depicting the asymmetric divisions of NBs throughout larval and pupal stages and recapitulating the above experiments. Scale bars, 20 μm.

### Single-cell RNA-seq of tNBs partly recapitulates the developmental trajectory

Our results indicate that Imp and Syp mediate tNB self-renewing potential and tumor hierarchy, at least partly, through the regulation of *chinmo*. We reasoned that this should lead to different transcriptional signatures in Imp^+^ and Syp^+^ tNBs that could help characterizing their molecular identity. We performed single-cell RNA-seq on dissociated and FACS-sorted GFP+ tNBs obtained from 56 adult *pox*>*pros*^*RNAi*^ tumors. Using the Chromium (10x Genomics) approach, we sequenced 5796 cells with a median number of 1806 genes/cell. As expected, NB identity genes like *miranda* (*mira*) or *deadpan* (*dpn*) appeared homogeneously expressed on the t-SNE plots while *elav* or *repo* that respectively mark neurons and glia are mostly absent (Figure 6B and Figure 6-figure supplement 1A). This indicates that the sequenced cells represent NB-like progenitors. *chinmo* was expressed in all cells, consistent with a translational control (Figure 6C). The unsupervised clustering method (graph-based) identified 8 sub-clusters (Figure 6A). Differential expression between the 8 sub-clusters uncovered *Imp* as the most up-regulated gene in one cluster of 468 cells (Cluster 7) relative to the rest of the dataset (Figure 6D and Figure 6-figure supplement 1B). In contrast, another cluster of 1203 cells (Cluster 1) was enriched in Syp (Figure 6-figure supplement 1B). However, Syp mRNA was also expressed at significant levels throughout all 8 clusters including Cluster 7 that contained Imp^+^ cells (Figure 6E). This indicates that Syp could be regulated at the translational level. Interestingly, Eip93F (E93) emerged as the most up-regulated gene in the rest of the tNB population when compared to the Imp^+^ Cluster 7 (Figure 6-figure supplement 1C). Moreover, when assessing the expression pattern of Imp and E93 on the t-SNE plot, we found that they were expressed in a complementary pattern that encompasses most cells (Figure 6D, F). This complementary pattern of expression is recapitulated at the protein level when staining tumors in adults with anti-Chinmo and anti-E93 antibodies (Figure 6G). Importantly, E93, like Syp, has recently been shown to be a marker for late NBs during development (Syed et al., 2017). This data thus shows that the populations of Imp^+^ tNBs and Syp^+^ tNBs can respectively be distinguished by Imp and E93 RNA levels.

During development, the transition from the Chinmo^+^Imp^+^ state to the Syp^+^E93^+^ state around early L3 requires the sequential expression of the temporal transcription factors *castor* (*cas*) and *seven-up* during the Chinmo^+^ Imp^+^ window (Maurange et al., 2008; Narbonne-Reveau et al., 2016; Ren et al., 2017; Syed et al., 2017). If this would still be the case in tumors then, *cas* and *seven-up* expression should be more significant in the Imp^+^ cluster 7. However, this was not the case (Figure 6-figure supplement 1D) indicating that Cas and Seven-up are not involved in the Chinmo^+^Imp^+^→Syp^+^E93^+^ transition in tNBs. This result is consistent with our previous observation that Seven-up is not detected by immunostaining in the tumor (Narbonne-Reveau et al., 2016). Moreover, the transcription factor *grainyhead* (*grh*) is not expressed in early NBs during embryogenesis and normally becomes transcriptionally activated in NBs at the end of embryogenesis. However, we find that *grh* is highly expressed in all if not all tNBs (Figure 6-figure supplement ID). This shows that tNBs, and in particular the CSC-like Imp^+^ tNBs, do not revert to an early embryonic-like state. Instead, Imp^+^ tNBs seem to have escaped regulation by temporal transcription factors and remain locked in an early larval identity.

To investigate whether tNBs in the tumor may form a continuum of differentiation states, we performed an unsupervised pseudotime analysis based on the variance in each gene’s expression across cells using Monocle (Trapnell et al., 2014) (Figure 6H). When assessing the relative expression of Imp and E93 with respect to Monocle ordering, we found that the states with highly expression of Imp were put at the beginning of the pseudotime (Figure 6I). Then a progressive down-regulation of Imp was observed along the pseudotime concomitant with an up-regulation of E93 that is then stabilized until the end of the pseudotime (Figure 6J). Thus, pseudotime analysis recapitulates a differentiation trajectory/hierarchy within tumors that is reminiscent to the differentiation trajectory observed in NBs during development. Altogether, these data indicate that the Imp/Syp-mediated tumor hierarchy can be characterized at the transcriptional level, and that the Chinmo^+^Imp^+^ → Syp^+^E93^+^ transition is not controlled by temporal transcription factors in the tumor context but by another unknown mechanism.

**Figure 6.**
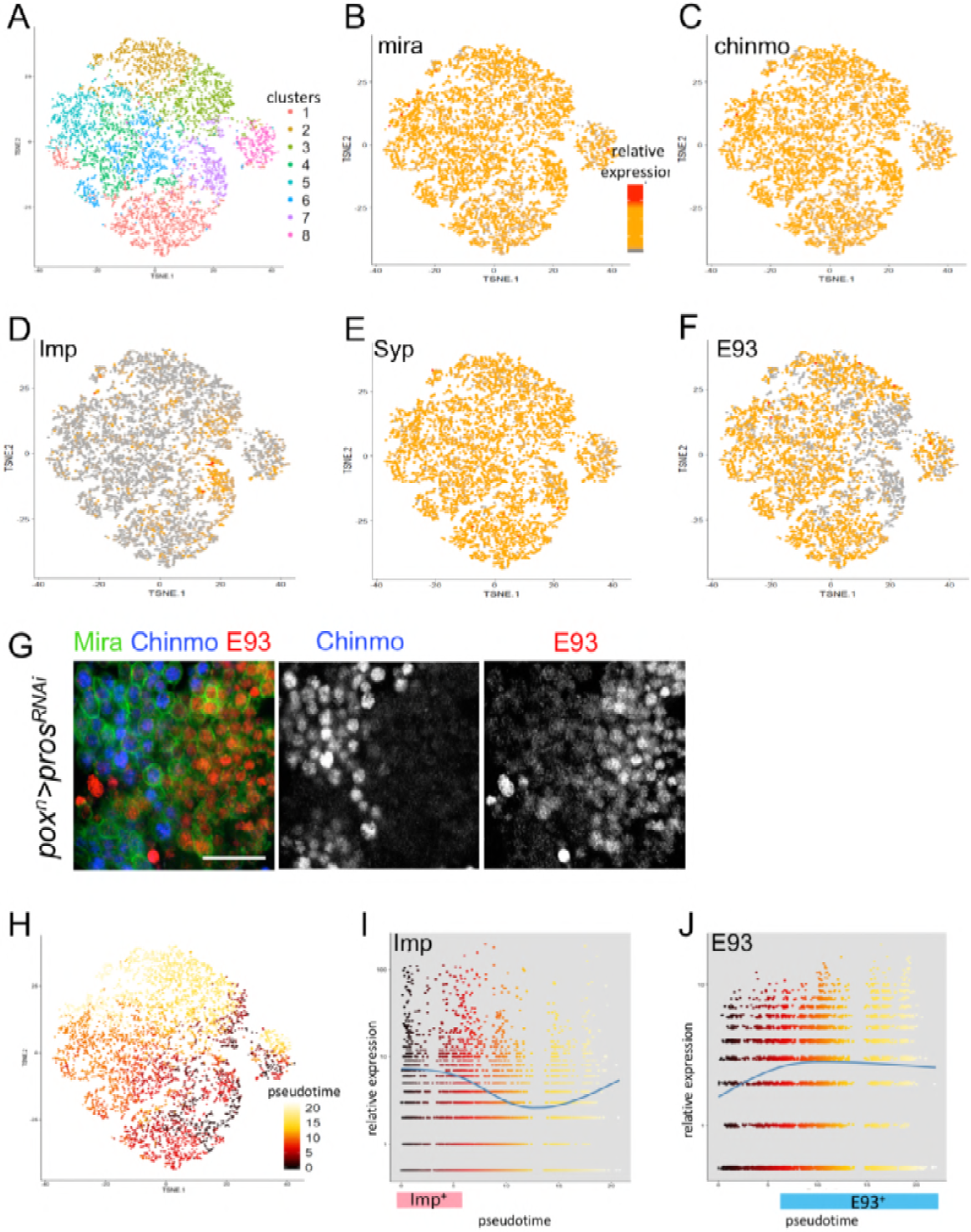
Single-cell RNA-seq and pseudotime analysis of tumors. (A) Unsupervised t-SNE plot generated by Cell Ranger using the graph-based clustering algorithm on sequenced tumor cells isolated by FACS. (**B,C,D,E,F**) Relative RNA levels of mira, chinmo Imp, Syp and E93. (**G**) E93 and Chinmo are expressed in a complementary pattern in adult tNBs (labelled with Mira). Scale bar 20μm. (**H**) Unsupervised pseudotime analysis by Monocole defines 20 states, color-coded on the t-SNE plot. (I) Expression of Imp with respect to pseudotime produced by Monocle. (**J**) Expression of E93 with respect to pseudotime produced by Monocle. Each color represents a pseudotime state.

### Syp triggers a metabolic switch

To investigate how Syp lowers the proliferation potential of tNBs, we performed RNA-seq on bulk *pox*^*n*^>*pros*^*RNAi*^ and *pox*^*n*^>*pros*^*RNAi*^ *Syp*^*RNAi*^ tumors (including the VNCs) of adult tumors. Between the two conditions, the expression of generic NB identity genes (*grh*, *dpn*, *ase*) did not significantly change (Figure 7A). In contrast, temporal NB markers such as *Imp*, *lin-28* and *E93* varied greatly, with early and late markers being respectively strongly enriched and down-regulated in the *pox*^*n*^>*pros*^*RNAi*^, *Syp*^*RNAi*^ tumors. This is consistent with our previous finding that Syp is required to silence *chinmo* and *Imp* and other early NB genes such as *lin-28* (Narbonne-Reveau et al., 2016; Syed et al., 2017). Interestingly, reactome pathway analysis identified the glycolytic and the ATP synthesis/TCA cycle/respiratory electron transport pathway as the most significantly enriched pathways in the *pox*^*n*^>*pros*^*RNAi*^, *Syp*^*RNAi*^ tumors (Figure 7A, B). Thus, loss of Syp in tNBs triggers an increase in the expression of glycolytic and respiratory genes in tumors, both pathways being well-known drivers of tumor growth (DeBerardinis and Chandel, 2016; Lunt and Vander Heiden, 2011; Wheaton et al., 2014). However, transcriptomic analysis on bulk tumors cannot discriminate whether expression of metabolic gene is increased in tNBs or in stromal cells (glia, neurons etc)-as a possible consequence of the reduced tumor heterogeneity. To resolve this question, we asked whether tNB metabolism evolved along the tNB trajectory reconstructed from the single-cell RNA-seq performed on*pox*^*n*^>*pros*^*RNAi*^ tumors. If Syp was regulating the metabolic state of tNBs, then a downregulation of glycolytic and respiratory should follow the Imp-to-Syp/E93 transition on the pseudotime. We found that glycolytic and respiratory/mitochondrial genes were indeed highly expressed in Imp^+^ tNBs (characterized by high levels of Imp mRNA) (Figure 7C) and many of them underwent a sharp down-regulation of their expression after the Imp→E93 transition. In agreement with a reduced glycolytic metabolism affecting cell growth after the transition, Syp^+^E93^+^tNBs exhibited a smaller size than Imp^+^ tNBs in tumors (Figure 7D and Figure 7-figure supplement 1). Moreover, along the pseudotime, the drop in metabolic genes correlated with that of cell cycle genes that were highest in tNBs around the Imp→E93 transition (Figure 7C). Cells located at the end of the pseudotime expressed very low levels of cell-cycle genes consistent with a significant proportion of Syp^+^E93^+^ tNBs being quiescent (about 26% in tumors of 2 days old adults based on our clonal analysis (See Material & Method). Together, these transcriptomic data demonstrate that Syp triggers a metabolic switch through the progressive silencing of glycolytic and respiratory/mitochondrial genes. This is likely to cause an exhaustion of the growth and self-renewing potential of both normal NBs and tNBs leading to their quiescence and/or terminal differentiation (Figure 7E).

## Discussion

### A subverted developmental transition triggers hierarchical tumors

We had previously shown that loss of Pros by itself is not sufficient to explain uncontrolled tumor growth. Indeed, *pros* inactivation in late larval NBs causes transient NB amplification but all NBs differentiate during metamorphosis. Instead, *pros* inactivation can cause unlimited tumor growth only if initiated in NBs (or GMCs) during early larval development when they are Chinmo/Imp^+^ (Narbonne-Reveau et al., 2016). Here we show that *pros* knock-down triggers the formation of an initial pool of Chinmo^+^Imp^+^ tNBs that proliferates but fail to systematically undergo the Imp→Syp transition that normally occurs in NBs during mid-larval development. Instead, a sub-population of Chinmo^+^Imp^+^ tNBs is permanently maintained (about 20-25% of tNBs) while the rest of tNBs are Syp^+^. Appearance of tumor heterogeneity from an initial homogenous population of Chinmo^+^Imp^+^ tNBs, population dynamics, and permanent tumor growth result from a division pattern in which Chinmo^+^Imp^+^ tNBs have a preference for symmetric self-renewing divisions (~64% of divisions) while also being able to generate Syp^+^ tNBs via asymmetric divisions (they self-renew while giving rise to a Syp^+^ tNB in ~15% of divisions), or symmetric differentiation divisions (they give rise to two Syp^+^ tNBs in ~21% of divisions). In addition, we have shown that Syp^+^ tNBs do not give rise to Chinmo^+^Imp^+^ tNBs and have a high tendency to enter quiescence. Given that the co-expression of Chinmo and Imp is sufficient to promote unlimited NB selfrenewal (Narbonne-Reveau et al., 2016), this division pattern is sufficient to promote unlimited NB amplification, creating hierarchical tumors with Chinmo^+^Imp^+^ tNBs acting as CSC-like cells while Syp^+^ tNBs behave as TAP-like cells (Figure 7E).

Thus, in this context, CSCs do not emerge as a consequence of multiple genetic lesions leading to the aberrant activation of oncogenic pathways, but they rather proceed from the unbalanced regulation of an early developmental transition (the Imp→Syp transition) that determines the switch from a default self-renewing to a differentiation-prone state. This indicates that CSCs may be molecularly extremely close to early stem cells, at least in tumors with developmental origins. Altogether, our work supports a tumor model where heterogeneity and hierarchical organization is “simply” induced by a forced switch from asymmetric-to-symmetric NB divisions and the subsequent failure to properly pass a developmental transition that normally restricts the self-renewing potential of stem cells. Our work is a striking demonstration of a growing organ that becomes “locked” into a developmental program leading to endless growth and tumorigenesis. We believe that similar processes are likely to explain the development of childhood cancers.

### Regulation of CSC division modes within tumors

The reasons underlying this unbalanced Imp→Syp transition are unclear. Temporal transcription factors, that schedule a systematic Imp→Syp transition in NBs during development, do not seem to be involved in this process in the tumorigenic context. Importantly, we have tested different types of tumors types caused by the aberrant amplification of NBs (following inactivation of either *brat*, *nerfin-1* or the mis-expression an hyper-active form of *aPKC*). Strikingly, they all exhibit a deregulated Imp→Syp transition (Narbonne-Reveau et al., 2016) and data not shown). Thus, the unbalanced Imp→Syp transition is not specifically instigated by the loss of Pros. Instead, it is possible that the Imp→Syp transition is being coopted by new signaling pathways that arise upon initial NB amplification and changes in the initial surrounding niche. Such pathways could interfere with the normal temporal mechanism regulating the switch.

While the division pattern we have described with our numerical model provides estimates of probabilities, it says nothing as to how these probabilities are biologically set within the tumor. A possible scenario is that cell fate determination upon division relies on signals received by immediate neighboring tumor cells, resulting in effective probabilities at the scale of the whole tumor. However, the model presented here has no spatial information. In this context, adding space to the model, and complementing our analysis of clone size and composition with an analysis of spatial distributions might provide useful insights on the mechanisms that instruct whether a CSC should self-renew or differentiate upon division.

### Syp triggers a metabolic switch in neural progenitors

Our transcriptomic analysis indicate that Syp induces a global down-regulation of glycolytic and respiratory genes in tNBs. Given that glycolysis and respiration are both important for cell growth and proliferation (DeBerardinis and Chandel, 2016; Lunt and Vander Heiden, 2011; Wheaton et al., 2014), their high expression in Chinmo^+^Imp^+^ tNBs is likely to sustain tNB growth and self-renewal, while their down-regulation in Syp^+^ tNBs is likely to contribute to their rapid entry into quiescence. Consistently, while the expression of cell-cycle genes is highest around the Imp→Syp/E93 transition on the pseudotime, it is followed by a rapid drop. Thus, newly generated Syp^+^ tNBs may first exhibit rapid cell divisions and then a sharp slow-down and quiescence entry possibly triggered by metabolic exhaustion. The rapid division time of newly generated Syp^+^ tNBs is consistent with our numerical model that predicts a slightly faster division time for proliferating Syp^+^ tNBs (1,3d) than for Chinmo^+^Imp^+^ tNBs (1,6d). However, in our model, Syp^+^ tNBs exhibit a strong probability to enter quiescence that is independent of the number of divisions they have undergone. Implementing parameters that take into account the history of Syp^+^ tNBs might help generating simulations closer to reality. Our observation that Syp induces a cell-autonomous metabolic switch in tNBs that presumably exhaust their growth and self-renewing ability implies that a similar switch may operate in NBs during development to limit their division and make them competent to differentiation. The transcription factors downstream to Syp that mediate this transcriptional switch remain to be identified. However, Chinmo could positively regulate glycolytic and respiratory genes in early NBs and CSC-like tNBs.

**Figure 7.**
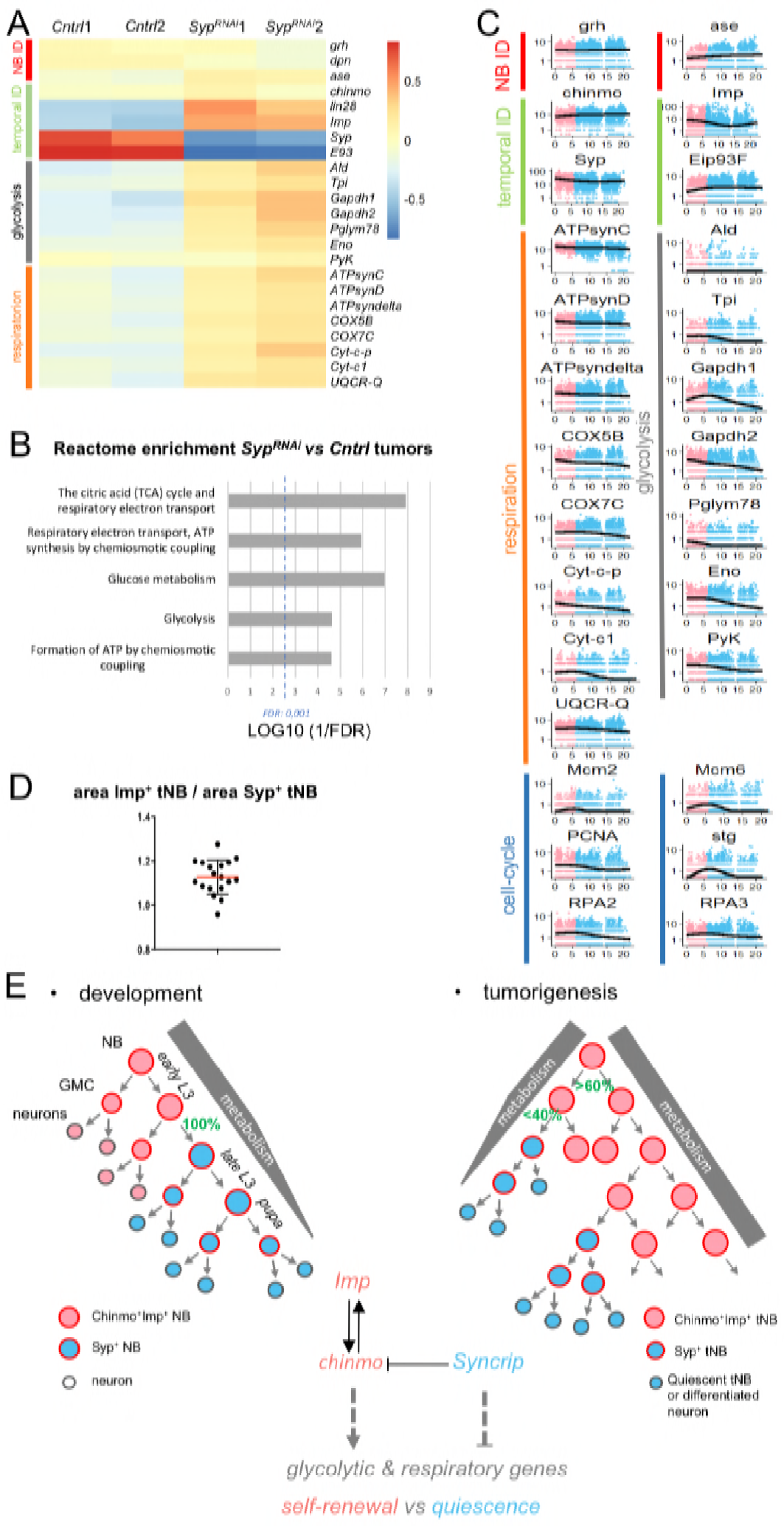
Syp regulates metabolic genes. (**A**) Heatmap of differentially expressed genes in the *pox*^*n*^>*pros*^*RNAi*^, *Syp*^*RNAi*^ vs *pox*^*n*^>*pros*^*RNAi*^ bulk tumors. (**B**) Enriched reactome pathways (*Panther*) from enriched genes in the *pox*^*n*^>*pros*^*RNAi*^, *Syp*^*RNAi*^ vs *pox*^*n*^>*pros*^*RNAi*^ comparison. (**C**) Expression of NB, temporal, glycolytic, respiratory and cell-cycle genes with respect to pseudotime produced by Monocle. (**D**) Each dot represents the average area of Imp^+^ tNBs over the average area of Syp^+^ tNBs for a single confocal section. (**E**) Antagonistic Imp and Syp regulate the self-renewing potential of NBs during development. Imp and Chinmo maintain a high metabolic activity to promote unlimited self-renewal, Syp silences metabolic genes to limit divisions and promote quiescence. Cooption of Imp and Syp in the tumor context establishes a tumor hierarchy characterized by a switch in cancer cell metabolism.

### Imp and Syp: a couple of antagonistic RBPs that defines CSC- and TAP-like identities

Chinmo and Imp are reminiscent to oncofetal genes in mammals, in that their expression decrease rapidly as development progresses while they are often mis-expressed in cancers. On these lines, the three IMP orthologs in humans (also called IGF2BP1-3) are also known as oncofetal genes. They emerge as important regulators of cell proliferation and metabolism in many types of cancers including pediatric cancers (Bell et al., 2015; Dai et al., 2017; Degrauwe et al., 2016a; Degrauwe et al., 2016b; Janiszewska et al., 2012). Along evolution, the ancestral *Syncrip* gene has been subjected to several rounds of duplication and has diverged into five paralogs in mammals, some of them emerging as tumor suppressors with an important role in tumor progression (Sakurai et al., 2016; Vanharanta et al., 2014).

Thus, the respective oncogenic and tumor suppressor roles of IMP and SYNCRIP gene families appear to have been conserved in humans and they may not be restricted to tumors of neural origin. Our study therefore raises the exciting possibility that these two families of RBPs may form a master module at the top of the self-renewal/differentiation cascades that regulate CSC populations and hierarchy in a wide spectrum of human cancers.

## Materials and Methods

### Fly culture

Drosophila stocks were maintained at 18°C on standard medium (8% cornmeal/8% yeast/1% agar).

### Fly lines

The protein trap line used was:

- Imp-GFP (Bloomington Stock Centre #G0080)

The GAL4 lines used were:

- *nab-GAL4* (Kyoto DGRC #6190) is a GAL4 trap inserted into *nab* (CG33545) that is active in all NBs of the VNC and central brain from late embryogenesis (Maurange et al., 2008).
- *pox*^*n*^-*GAL4* (Boll and Noll, 2002) is active in six thoracic NBs of the VNC.

The UAS lines used were:

- *UAS-Flybow. 1.1* (Bloomington Stock Centre #35537) (Hadjieconomou et al., 2011).
- *UAS-mCherry*^*chinmoUTRs*^ (Dillard et al.,2018). In NB tumors carrying *UAS-mCherry*^*chinmoUTRs*^, expression of *cherry* reflects the post-transcriptional regulation of endogenous *chinmo*.Consistently, we observe that Cherry always overlaps with endogeneous Chinmo as detected by immunostaining (Figure S4B).
- *UAS-SypF*^*RNAi1*^ (Vienna Drosophila RNAi Center #33011)
- *UAS-SypF*^*RNAi2*^(Vienna Drosophila RNAi Center #33012)
- *UAS-Syp-RB-HA* (T.Lee, Janelia Research Campus, Virginia, USA) (Liu et al., 2015).
- *UAS-pros*^*RNAi1*^ (Transgenic RNAi Project #JF02308, Bloomington Stock Centre #26745)
- *UAS-pros*^*RNAi2*^ (Vienna Drosophila RNAi Center #101477)
- *UAS-Imp*^*RNAi*^ (Vienna Drosophila RNAi Center #20322)
- *UAS-chinmo*^*RNAi*^ (Transgenic RNAi Project #HMS00036, Bloomington Stock Centre #33638)
- *UAS-dicer2* (Bloomington Stock Centre #24650 and #24651) was used in combination with GAL4 lines in order to improve RNAi efficiency.
- *UAS-mCD8::GFP* (Bloomington Stock Centre #5130 and #32185) and *UAS-myr::GFP* (Bloomington Stock Centre *#32197*) were used to follow the driver expression.
- *UAS-mCherry.NLS* (Bloomington Stock Centre #38424)
- *UAS-Imp*^*RNAi*^; *UAS-Syp*^*RNAi2*^(*UAS-Syp*^*RNAi2*^ from Vienna Drosophila RNAi Center #33012)
- *UAS-Syp*^*RNAi*^; *UAS-chinmo*^*RNAi*^ (*UAS-Syp*^*RNAi1*^ from Vienna Drosophila RNAi Center #33011)
- *UAS-pros*^*RNAi1*^, *UAS-mCD8::GFP* (*UAS-pros*^*RNAi1*^ from Transgenic RNAi Project #JF02308)
- *UAS-pros*^*RNAi2*^; *UAS-mCherry*^chinmoUTRs^(*UAS-pros*^*RNAi2*^ Vienna Drosophila RNAi Center #101477)

Fly crosses were set up and raised at 29°C. To label all tumor cells with GFP, *pox*^*n*^-*GAL4*, *UAS-pros*^*RNAi*^, *UAS-dicer2 flies* were crossed with the G-TRACE system *UAS-FLP*, *Ubi-p63EFRTstopFRTnEGFP* (Bloomington Stock Centre #28282) (Evans et al., 2009). To generate RFP+ clones in tumors, *pox*^*n*^-*GAL4*, *UAS-pros*^*RNAi1*^, *UAS-mCD8::GFP*; *UAS-dicer2* flies were crossed to *UAS-Flybow.1.1; hs-mFLP5* (Bloomington Stock Centre #35534 and #35537) (Hadjieconomou et al., 2011). The progeny of this cross was collected 48 hours after adult eclosion and heat-shocked for 1h30’ at 37°C. Flies were put back at 29°C after heat-shock. Note that differentiated neurons generated from RFP+ tNBs will not retain RFP expression due to the silencing of*pox*^*n*^-*GAL4* in neurons.

### Image processing

Confocal images were acquired on a Zeiss LSM780 microscope. FIJI-ImageJ was used to process confocal data and to compile area and volume data. Imaris Image Analysis Software (http://bitplane.com) was used to generate 3D representations of clones in tumors.

### Tumor segmentation using Tissue Analyzer

For figures 2F and S6, *pox*^*n*^>*pros*^*RNAi*^ tumors from 2 to 5 day-old adult flies carrying an Imp-GFP marker were dissected, fixed in 4% paraformaldehyde and stained with antibodies against Miranda and GFP proteins. VNCs were visualized in a Zeiss LSM780 confocal microscope and co-planar images were made of several sections ranging from the tumor surface to its interior. Several confocal planes from each 4 tumor (n = 21) were analyzed using the “Tissue Analyzer” plugin for FIJI-ImageJ (Aigouy et al., 2016). This tool allows for cell membrane tracing, via the signal from Miranda which marks the cellular border of all tumor cells, segmentation and quantification of several parameters, including size, of each tumor cell individually. CIL^+^ tNBs are identified by their strong Imp-GFP signal. Initial automatic segmentation was further refined manually to ensure all cell boundaries were properly tracked and cells assigned a correct identity. Data points represent different planar sections within the same tumor, in order to account for variability within each tumor.

### Statistical analysis

For each experiment, at least 3 biological replicates were performed. Biological replicates are defined as replicates of the same experiment with flies being generated by different parent flies. For all experiments, we performed a Mann-Whitney test for statistical analysis. Statistical analysis was performed using GraphPad Prism version 7.04 for Windows (GraphPad Software, La Jolla California USA, http://www.graphpad.com). Results are presented as dot plots. Error bars represent s.d. from the mean. The sample size (n) and the *P*-value are reported in the figure legends (\*\*\*\**P*≤0.0001; \*\*\**P* ≤0.001; \*\**P* ≤0.01; \**P*≤0.05; ns, not significant).

### Immunohistochemistry

Larval and adult VNCs were dissected in phosphate-buffered saline (PBS) and fixed for 10 min at room temperature (RT) in 4% paraformaldehyde/PBS. VNCs were rinsed in PBT (PBS containing 0.5% Triton X-100) and incubated with primary antibody overnight at 4°C. Secondary antibodies (Jackson ImmunoResearch) were incubated overnight at 4°C. VNCs were mounted in Vectashield (Clinisciences, France) with or without DAPI for image acquisition. The following primary antibodies were used: chicken anti-GFP (1:1000, Aves #GFP-1020), rabbit anti-RFP (1:500, Rockland #600-401-379), rat anti-RFP (1:500, Chromotek #5F8), mouse anti-Miranda (1:50, A. Gould, Francis Crick Institute, London, UK), rabbit anti-PH3 (1:500, Millipore #06-570), rat anti-PH3 (1:500, Abcam #AB10543), rat anti-Elav (1:50, DSHB #9F8A9), rabbit anti-cleaved Dcp-1 (1:500, Cell Signaling #9578)), rat anti-Chinmo (1:500, N. Sokol, Indiana University, Bloomington, USA), rabbit anti-Imp (1:500, P. Macdonald), guinea pig anti-Syp (1/500, I. Davis, Oxford University, UK), rabbit anti-Syp (1/200, I. Davis, Oxford University, UK), rabbit anti-E93 (1/2500 D.J. McKay, University of North Carolina, Chapel Hill).

### Affinity pull-down assays

Affinity pull-down assays were performed as previously described (Medioni et al., 2014), and Western Blots performed using rat anti-Imp and rabbit anti-Syp antibodies. The 5’ and 3’ UTRs of chinmo were cloned from the cDNA clone #RE59755 (DGRC, EST Collection) using the following primers:

Chin-5′UTR/pB S_Forward **aatttctagaAGTCAAAAAGAAACTGCCGTG**

Chin-5′UTR/pBS_Reverse gatg**aagctt**GGTGCCAGCAGTGATGCT

Chin-3′UTR/pB S_Forward taaa**tctaga**GAAGCAGCCGCAACAGCA

Chin-3′UTR/pB S_Reverse tttt**aagctt**GGTGAATTTTCATTTGTACGAAGAA

Capitals represents analogous or complementary parts of the chinmo sequence. In bold are the restriction sites used for cloned into pBS-KS(-) (TCTAGA=*Xba*I, AAGCTT=*Hind*III), and in lower case are extra bases for efficient digestion. This plasmid was used to generate the UTP-biotinylated-RNA probes (5’UTR and 3’UTR).

### RNA extraction of bulk tumors

To generate NB tumors *pox*^*n*^-*GAL4*, *UAS-mCherry.NLS*; *UAS-dicer2 flies* were crossed to flies carrying:

1. *UAS-pros*^*RNAi1*^
2. *UAS-Syp*^*RNAi1*^; *UAS-pros*^*RNAi1*^

Crosses were grown at 29°C. 16 adult females and 13 adult males (8 day-old) were killed in 70 % ethanol and washed once in PBS. VNCs were dissected in PBS and collected in a RNase-free Protein LoBinding tube filled with 750μL of Lysis Buffer (Buffer RLT, RNeasy Mini Kit, Qiagen) supplemented with 7.5μL of β-mercaptoethanol and 500μL of glass beads (diameter 0.75-1mm, Roth, A554.1). Samples were disrupted and homogenized using the tissue homogenizer Precellys 24 (Bertin Technologies).

Sample tubes were then stored at −80°C up to RNA extraction. Total RNA was extracted using the RNeasy Mini Kit (Qiagen). RNA quality and quantity were checked by running samples on an Agilent RNA 6000 Pico Chip (Agilent Technologies). Biological triplicates were made for each condition: (1) n=29, n=29, n=29; (2) n= 29, n=29, n=29.

### Library preparation and sequencing of bulk tumors

Total RNA-Seq libraries were generated from 130 ng of total RNA using TruSeq Stranded Total RNA LT Sample Prep Kit with Ribo-Zero Gold (Illumina, San Diego, CA), according to manufacturer’s instructions. Briefly, cytoplasmic and mitochondrial ribosomal RNA (rRNA) was removed using biotinylated, target-specific oligos combined with Ribo-Zero rRNA removal beads. Following purification, the depleted RNA was fragmented into small pieces using divalent cations at 94°C for 2 minutes. Cleaved RNA fragments were then copied into first strand cDNA using reverse transcriptase and random primers followed by second strand cDNA synthesis using DNA Polymerase I and RNase H. Strand specificity was achieved by replacing dTTP with dUTP during second strand synthesis. The double stranded cDNA fragments were blunted using T4 DNA polymerase, Klenow DNA polymerase and T4 PNK. A single ‘A’ nucleotide was added to the 3′ ends of the blunt DNA fragments using a Klenow fragment (3′ to 5′exo minus) enzyme. The cDNA fragments were ligated to double stranded adapters using T4 DNA Ligase. The ligated products were enriched by PCR amplification (30 sec at 98°C; [10 sec at 98oC, 30 sec at 60oC, 30 sec at 72°C] x 12 cycles; 5 min at 72°C). Surplus PCR primers were further removed by purification using AMPure XP beads (Beckman-Coulter, Villepinte, France) and the final cDNA libraries were checked for quality and quantified using capillary electrophoresis.

The libraries were loaded in the flowcell at a concentration of 2.8 nM and clusters were generated using the Cbot and sequenced on an Illumina HiSeq 4000 system as paired-end 100 base reads following Illumina’s instructions.

### RNA-Seq analysis of bulk tumors

Quality control of raw reads was done using FastQC. Reads were mapped to *dmel6* reference genome with *Subread aligner* using default parameters (Liao et al., 2013). Read counts were calculated using *Subread featureCount* (Liao et al., 2014). Differential expression was computed using R package DESeq2 (Love et al., 2014). To concentrate on genes that are expressed in tNBs and remove those that are specific to the stroma or VNC neurons, we excluded from the bulk RNA-seq analysis all genes that are not expressed in at least 0,5% of the 5796 cells used for the single-cell RNA-seq experiment. Among the remaining 6660 genes, we excluded genes with a basemean <1000 and selected as significantly enriched genes those with an adjusted p-value <0,001. Gene Ontology and Reactome Pathway analysis was made using *Panther* (http://www.pantherdb.org/).

### Preparation of tNBs for single-cell mRNA-seq

NB tumors were generated in *pox*^*n*^-*Gal4*, *UAS-GFP UAS-pros*^*RNAi2*^, *UAS-dicer2* flies. Fifty-six adult females (6-8 day-old) were killed in 70 % ethanol and washed once in PBS. VNCs were dissected in PBS, collected in a RNase-free Protein LoBinding tube filled with ice-cold PBS, and incubated in a freshly prepared dissociation solution containing 0,4 % bovine serum albumin (BSA), 1 mg/mL collagenase I and papain (Sigma Aldrich) in PBS for 1 h 15 min at 29 °C with lowest agitation. Tissues were then disrupted manually by pipetting up and down with a 200 μL tip. Dissociated cells were pelleted for 20 min at 300 g at 4 °C to remove the dissociation solution and to resuspend cells in ice-cold PBS + 0,4 % BSA. The cell suspension was filtered through a 30 μm mesh Pre-Separation Filter (Miltenyi) to remove debris and transferred in new RNase-free Protein LoBinding tube for FACS sorting. Forty thousand GFP^+^ tNBs cells were isolated using a FACSAriaII machine (BD) with a 85 μm nozzle, at 45 psi low pressure and according to viability, cell size and GFP intensity. In the next 30 min, sorted-cells were encapsidated using with the Single Cell Controller (10X genomics) for single-cell RNA-seq

### Single-cell mRNA sequencing & analysis

#### Single-cell mRNA sequencing

Single cells were processed using the Single cell 3’ Library, Gel beads & multiplex kit (10X Genomics, Pleasanton) as per the manufacter’s protocol. Cells, were partitioned into nanoliter-scale Gel Bead-In-Emulsions (GEMs) with the Chromium Single Cell Controller (10X genomics, Pleasanton), where all generated cDNA share a common 10x Barcode. Libraries were generated and sequenced from the cDNA and the 10x Barcodes are used to associate individual reads back to the individual partitions. Analysis using molecular indexing information provides an absolute digital measurement of gene expression levels. Sequencing was performed using a NextSeq 500 Illumina device (1sample) containing transcript lengh of 57 bp.

#### Data Processing

NextSeq data were analysed using the 10x Genomics suite Cell Ranger 2.0.1 and specifically the function mkfastq. This first level of analysis generates quality metrics (Q30, number of reads by sample…) and FASTQ files. Then Cell Ranger count is performed on the library. This second level of analysis consisted in mapping with STAR against drosophila reference genome, detecting the number of cells and genes per cell and finally quantifying gene expression (count tables) and performing t-SNE analysis.

#### Cell trajectory and pseudo time analysis by Single-cell mRNA sequencing

Bioinformatics analysis was performed using R workspace, *CellRanger R Kit* and *Monocle* version 2 (Qiu et al., 2017a; Qiu et al., 2017b; Trapnell et al., 2014). Graphics have been generated through *ggplot2* R library. *Cell Ranger R Kit* was used to build the CellDataSet required as *Monocle*’s input. UMI Counts were both normalized across cells using *estimateSizeFactors* function and gene expression variance was estimated using *estimateDispersions* functions to remove outlier genes using DESeq2 packages (Love et al., 2014). Marker genes were selected for subsequent analysis based on their normalized average expression and variability across cells using *dispersionTable* function. Dimension reduction was performed with *Monocle* implementation of Discriminative Dimension Reduction Tree (DDRTree) algorithm (Mao et al., 2017) using *reduceDimension* function. Minimum spanning tree computation, cell state assignations and pseudo time reconstruction was performed using *orderCells* functions.

### Numerical model

#### 1. Algorithm of the clone model

In this section we detail the stochastic numerical model used to generate clones.

Each clone grows from a single initial cell. There are two cell types, Chinmo^+^Imp^+^ tNBs (which we will call C cells for simplicity) and Syp^+^ tNBs (S). The probability that the initial cell is a C cell is *p*_*c*_ and the probability that the initial cell is a S cell is *p*_*s*_ = 1 − *p*_*c*_.

We assume that C and S cells have constant division rates, *k*_*c*_ and *k*_*s*_ respectively. We define the division times as the inverses of the division rates, so that *T*_*c*_ = 1/*k*_*c*_ and *T*_*s*_ = 1/*k*_*s*_. An implicit assumption is that we neglect any refractory period following a division. Divisions are considered as random events occurring at a certain rate.

We use T=1h long time steps. The growth of a model clone during 10 days thus runs over 240 time steps. At each time step, for each C cell (resp. S cell) in the clone, the probability to undergo division is equal to 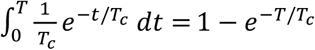.

If a C cell undergoes division, it can either:

- Generate 2 C cells, with probability *P*(*C* → *CC*)
- Generate 1 C cell and 1 S cell, with probability *P*(*C* → *CS*)
- Generate 2 S cells, with probability *P*(*C* → *SS*) = 1 - *P*(*C* → *CC*) - *P*(*C* → *CS*)

Similarly, if a S cell undergoes division, it can either:

- Generate 2 S cells, with probability *P*(*S* → *SS*)
- Generate 1 C cell and 1 S cell, with probability *P*(*S* → *CS*)
- Generate 2 C cells, with probability *P*(*S* → *CC*) = 1 - *P*(*S* → *SS*) - *P*(*S* → *CS*)

In addition, each newly created C cell (resp. S cell) has a probability *q*_*c*_ (resp. *q*_*s*_) to be quiescent. Note that this also applies to the initial cell of the clone. Quiescent cells lose their ability to undergo further divisions. In this simplified model, this probability does not depend on cell generation (as cells have no age in the model). However, it is likely that “older” cells have a higher chance to become quiescent. This is not taken into account here.

To produce plots of size distributions and of composition ratios, we simply generate a large amount of clones (N=1000) by iterating this algorithm. Due to the stochastic nature of the algorithm, there is important size and composition variability among simulated clones, which we can compare to the variability observed *in vivo*.

#### 2. Reducing the amount of independent parameters

This general model has 9 independent parameters: *p*_*c*_, *T*_*c*_, *T*_*s*_, *P*(*C* → *CC*), *P*(*C* → *CS*), *P*(*S* → *SS*), *P*(*S* → *CS*), *q*_*c*_ and *q*_*s*_.

To reduce the amount of independent parameters, we performed experimental measurements in clones that allow to establish quantitative relationships between the parameters.

##### a. 2-cell clones

First, we analyzed clones composed of exactly two cells. To that end, we chose to look tumors 2 days after clonal induction, as at this stage many clones are composed of two cells. The probability for a 2 cell clone to be composed of two C cells is:

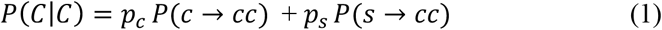

Similarly,

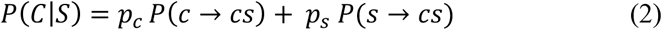

and

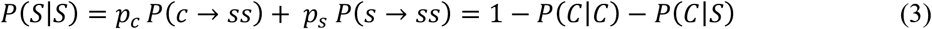

We estimated *P*(*C*|*C*), *P*(*C*|*S*) and *P*(*S*|*S*) from N=122 2-cell clones (7 animals), and found that *P*(*C*|*C*) ≈ 0.29, *P*(*C*|*S*) ≈ 0.07 and *P*(*S*|*S*) ≈ 0.64.

Using these values in equations (1) and (2) reduces the amount of independent parameters from 9 to 7 (as equation (3) is not independent from equations (1) and (2)).

##### b. Quiescent cells

We analyzed clones 8 days ACI. We reasoned that clones still composed of a single cell at this stage were most likely initiated from an initially quiescent cell. Hence the probability *P*(*QC*) to observe a clone composed of a single C cell at 8 days is equal to the probability for an initial cell to be a quiescent C cell, so that:

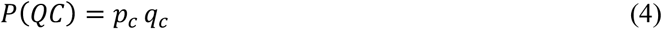

Similarly, the probability that a clone is composed of a single quiescent S cell is

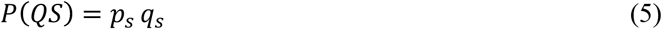

We estimated *P*(*QC*) and *P*(*QS*) from N=338 clones 8 days after induction (5 animals), and found that *P*(*QC*) ≈ 0.02 and (*QS*) ≈ 0.26. Using these values in equations (4) and (5) reduces the amount of independent parameters by another 2.

##### c. Hierarchical scenario

We are left with 5 independent parameters. We showed in the main text that experimental clone data has the signature of a hierarchical growth mode (maintenance of clones composed of S cells only). In the following we thus rule out the generation of C cells by S cells. Hence (*S* → *SS*) = 1, and *P*(*S* → *CS*) = *P*(*S* → *CC*) = 0

In addition, if C cells can only be generated by C cells, we can estimate the division time *T*_*c*_ of C cells. At time t, the number of cells in clones only composed of C cells should be on average equal to *N*_*c*_ (*t*) = *e*^*t*/*T*_*c*_^.

At 2 days, we find that clones composed only of C cells are composed, on average, of 3.4 cells (N=75 clones from 5 animals). Hence 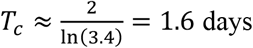.

Hence, we are left with only two independent parameters, from which all the others can be determined. We arbitrarily chose *T*_*s*_ and *P*(*C* → *CC*) as these two parameters, and used these as free parameters to perform the fits.

#### 3. Fitting experimental data

To determine the values of *T*_*s*_ and *P*(*C* → *CC*), we adopted a fitting approach. We want to determine the values of *T*_*s*_ and *P*(*C* → *CC*) that fit best our experimental data. To that end we generated error maps in the *P*(*C* → *CC*), *T*_*s*_) space (Figure SXXX). To generate the error maps, we computed 1000 clones for each pixel (100×100) on the map.

We calculate the error on size as the Kolmogorov-Smirnov distance between model and experimental clone size distributions, for all clone categories and at all stages (12 terms).

We calculate the error on composition as the normalized sum of Euclidian distances between model and experimental proportions of each clone category at all stages (12 terms).

The combined error is the average between the normalized error on size and the normalized error on composition.

We find that the smallest error is found for *P*(*C* → *CC*) ≈ 0.64 and *T*_*s*_ ≈ 1.3 days, and thus (Equations 1–5):

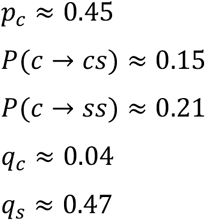

We use these values to generate Figures 3D-F and 4D, and Figure 3-figure supplement 1.

#### 4. Tumor composition and deterministic version of the model

##### a. Proportion of S and C cells in tumors

To test whether our model also accounts for the stabilization of the volume fraction occupied by S and C cells in tumor, we simulated the growth of clones on longer periods (30 days), and plotted the overall proportion of each cell type as a function of time (Figure 3F).

##### b. Deterministic, continuous model

To better understand this homeostatic behavior without systematically simulating thousands of clones, we wrote a deterministic version of the model where the exact same ingredients and parameters are used.

For simplicity, we call *C* the number of C cells, *S* the number of *S* cells, *C*_*q*_ the number of quiescent C cells and *S_q_* the number of quiescent S cells. The rate of change of *C* writes:

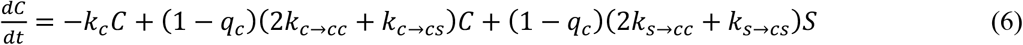

With *k*_*x* → *yz*_ = *k*_*x*_*P*(*x* → *yz*), where *x,y* and *z* are cell types (C or S)

Each C cell division removes 1 C cell (first term). C cell divisions can also generate 1 or 2 C cells with given rates (second term), with a possibility that they are quiescent (prefactor (1 − *q*_*c*_)). S cell divisions can also generate C cells (third term), which can also be quiescent (prefactor).

Similarly, we can write the rate of change of *S*:

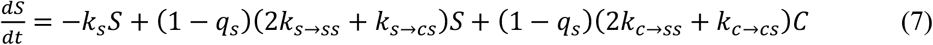

And for quiescent cells:

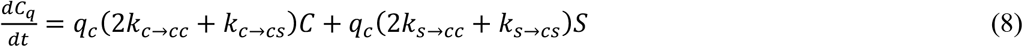

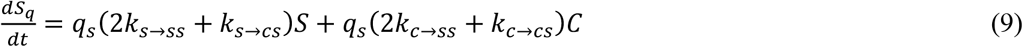

Note that this continuous model is fully deterministic, and therefore not suited for the simulation of clone size distribution and composition, which are inherent to the stochastic aspects of divisions. However it can recapitulate the overall proportion of cell types in a large population of cells. Integrating this system of equations, we thus computed the ratios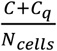 and 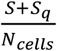, that is, the respective proportions of C and S cells. We first compared the outcome of the deterministic model to the outcome of the clone model using the exact same parameters. We find that the deterministic model perfectly fits the stochastic model (Figure 3F), and reaches the same homeostatic state. Importantly, this is also a control that both algorithms were properly implemented. We also use this deterministic model to illustrate the influence of the quiescence probability *q*_*s*_ (Figure 3-figure supplement 2).

## Acknowledgements

We thank I. Davis, P. Macdonald, D.J. McKay, C. Meignin M. Noll, N. Sokol for flies and antibodies. We are grateful to B. Aigouy for helping with the “Tissue Analyser” plugin, A. Saurin for alignment of RNA-seq data, F. Hubert and J. Oliver for discussions on mathematical modeling, J.C Patel for help with image analysis and P. Grenot for help with FACS. We also acknowledge the Bloomington Drosophila Stock Center (NIH P40OD018537), the Vienna Drosophila RNAi Center (VDRC), TRiP at Harvard Medical School (NIH/NIGMS R01-GM084947), Kyoto DGRC and NIG-Fly Stock Centers, and the Developmental Studies Hybridoma Bank (DSHB) for monoclonal antibodies. Bulk tumor sequencing was performed by the IGBMC GenomEast platform, a member of the ‘France Génomique’ consortium (ANR-10-INBS-0009)”. We thank France-BioImaging/PICsL infrastructure (ANR-10-INSB-04-01). We thank S. Kerridge for critical reading of the manuscript.

## Supplementary figures

**Figure 1 figure supplement 1:**
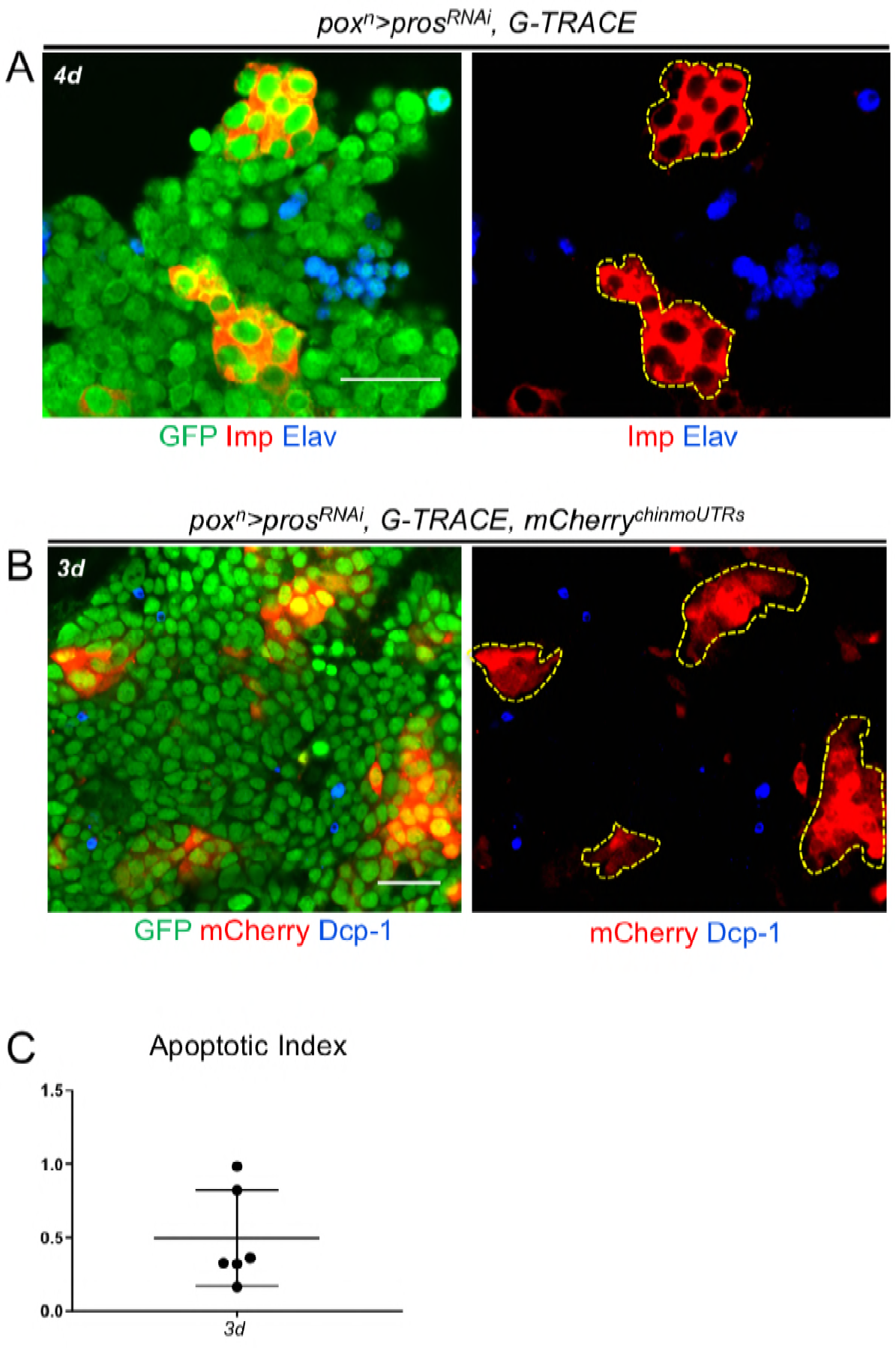
Composition of tumors that persist in adults. (**A**) Proportion of Elav+ cells in 1-day-old (n=7 VNC) and 4-day-old (n=7 VNC) *pox*^*n*^>*pros*^*RNAi*^ tumors. *P*= 0.165. (**B**) Volume of Elav+ cells in in 1-day-old (n=7 VNC) and 4-day-old (n=7 VNC) *pox*^*n*^>*pros*^*RNAi*^ tumors. *P*=0.004. (**C**) Neurons, marked with Elav, are dispersed outside the CIL+ clusters in 4-day-old *pox*^*n*^>*pros*^*RNAi*^ tumors. tNBs are marked with GFP. CIL+ tNBs are identified by the presence of Imp and delimited by dashed lines. (**D**) Apoptotic cells, marked with Dcp-1, are rare in 3-day-old *pox*^*n*^>*pros*^*RNAi*^ tumors and mostly found among CIL^-^ negative tNBs. tNBs are marked with GFP. The *UAS-mCherry*^*chinmoUTRs*^ is expressed in tumors where it is post-transcriptionally regulated to reflect *chinmo* expression (Fig. S4B). CIL+ tNBs are thus identified by the presence of mCherry and delineated by dashed lines (**E**) Apoptotic index in 3-day-old *pox*^*n*^>*pros*^*RNAi*^ tumors (n=6 VNC). Scale bars, 20 μm.

**Figure 3-figure supplement 1:**
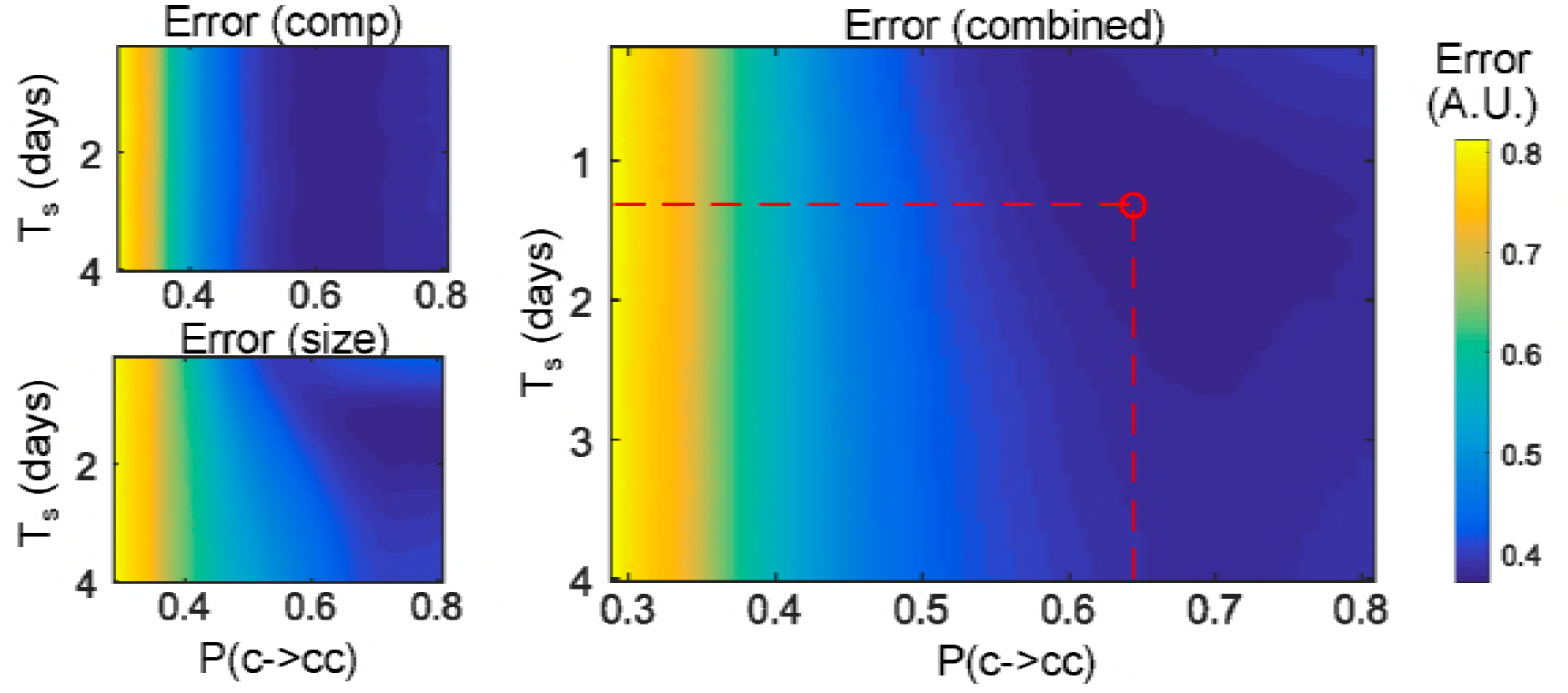
Error maps. Normalized error maps in the (Ts,P(c->cc)) space. Top left: error on clone compositions; bottom left: error on clone sizes; right: combined error.

**Figure 3-figure supplement 2:**
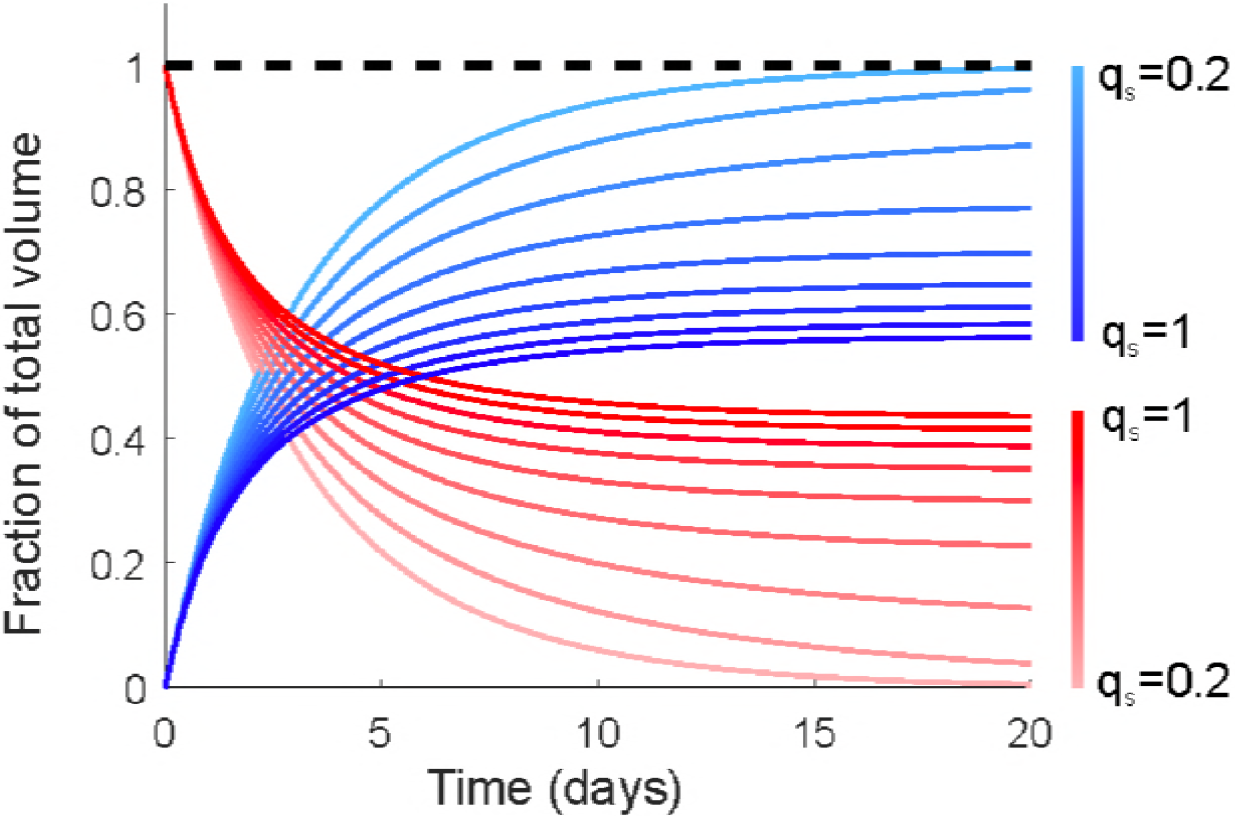
Impact of Syp^+^ tNB quiescence probability on tNB proportion. Proportion of Syp^+^ (blue) vs CIL^+^ (red) tNBs as predicted by the deterministic model for a wide range (20% to 100%) of Syp^+^ tNB quiescence probability.

**Figure 4 - figure supplement 1:**
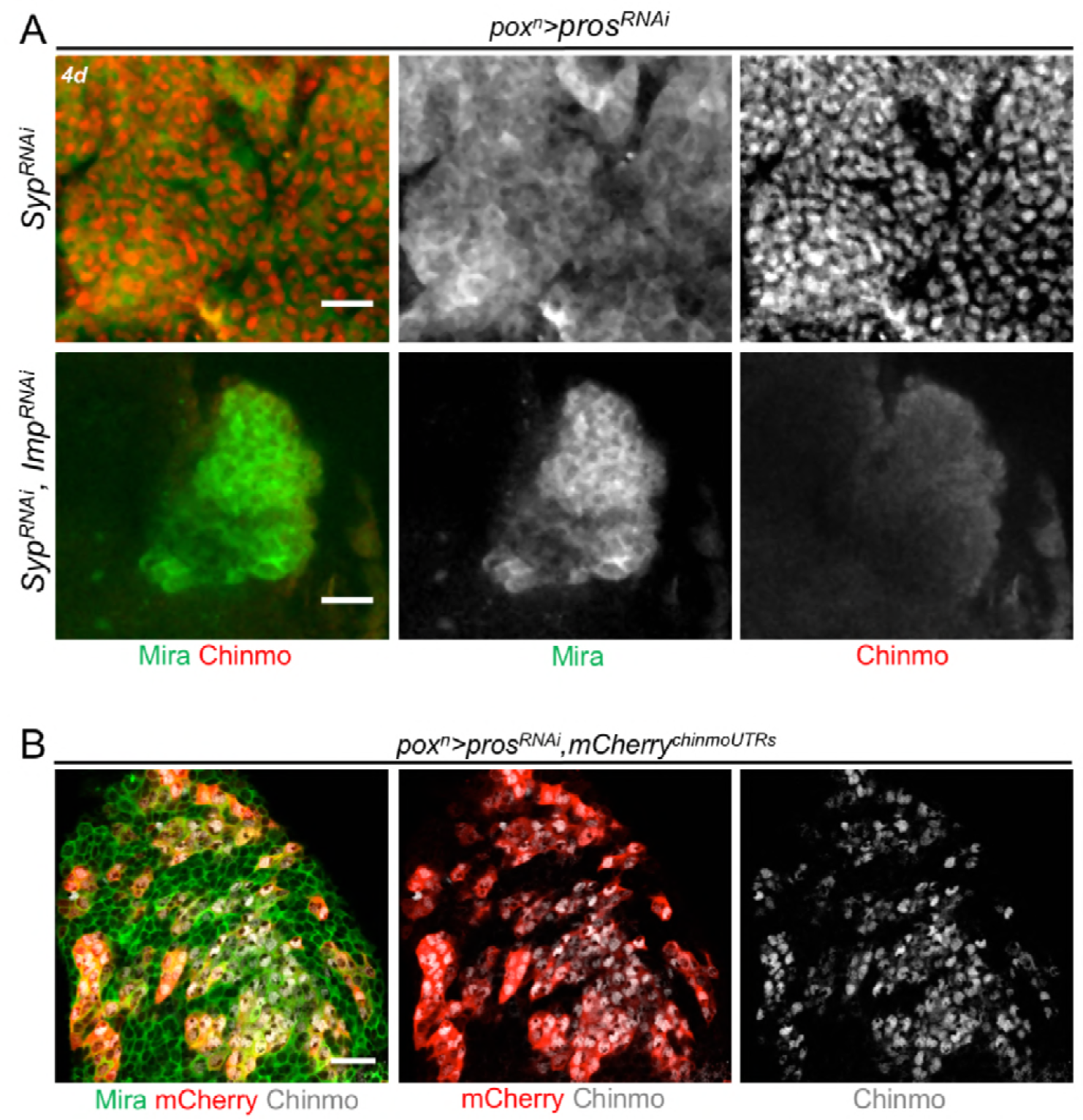
Syp and Imp antagonistically regulate Chinmo in tumors. (**A**) *pox*^*n*^>*pros*^*RNAi*^, *Syp*^*RNAi*^ tumors are mostly composed by Chinmo^+^ tNBs. Chinmo is silenced in *pox*^*n*^>*pros*^*RNAi*^, *Syp*^*RNAi*^, *Imp*^*RNAi*^ tumors indicating that Imp is necessary to promote Chinmo expression in *pox*^*n*^>*pros*^*RNAi*^, *Syp*^*RNAi*^ tumors. tNBs are marked with Mira. (**B**) The *UAS-mCherry*^*chinmoUTRs*^ transgene reflects endogeneous *chinmo* expression when expressed in tumors. Scale bars, 20 μm.

**Figure 6 - figure supplement 1:**
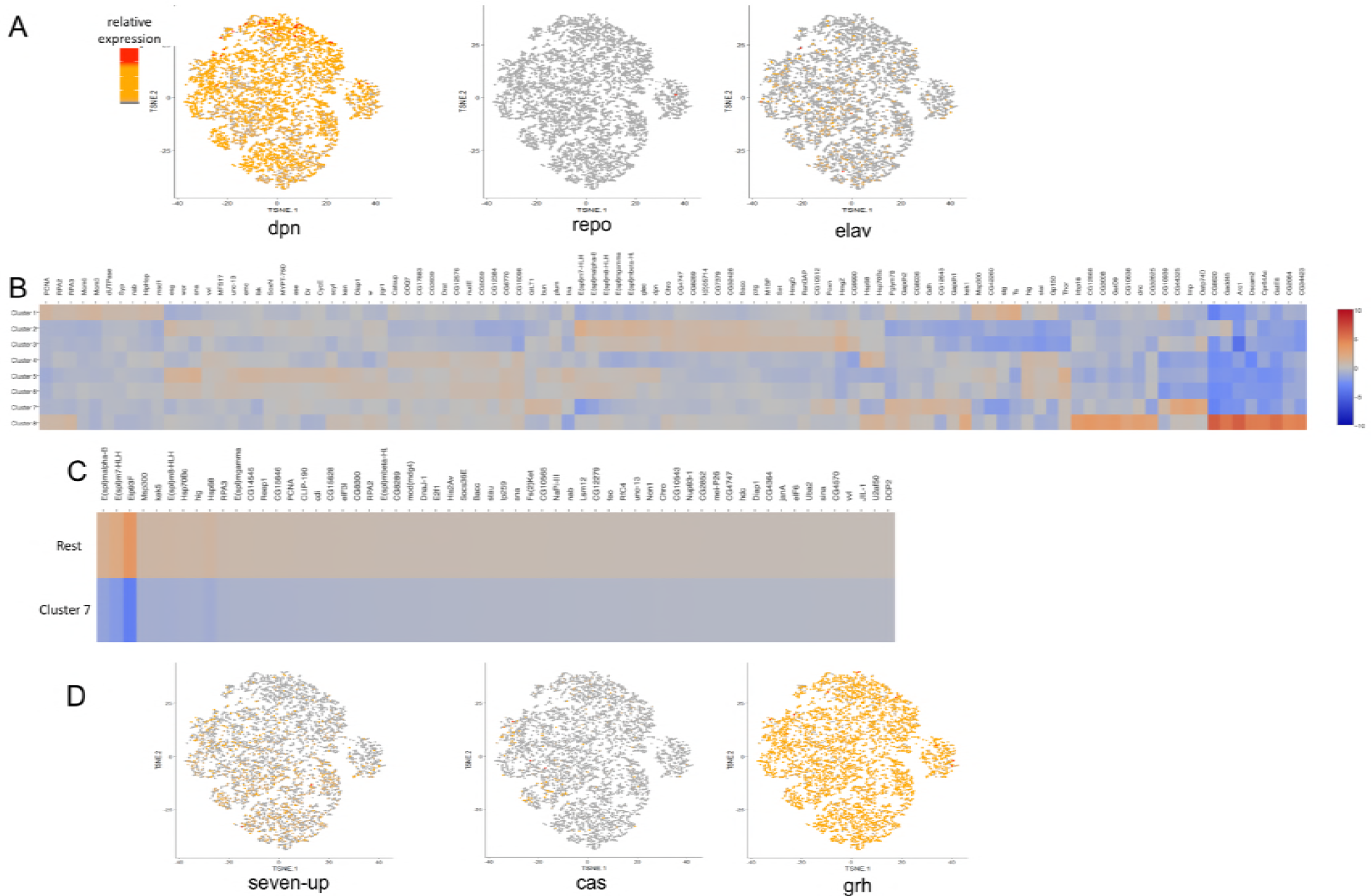
Identification of differentially expressed genes by single-cell RNA-seq. (**A**) Sequenced cells express typical NB identity markers such as dpn, while glial (repo) or neuronal (elav) genes are not expressed. (**B**) Heatmap depicting the top-upregulated genes in clusters 1 to 8. (**C**) Heatmap depicting the top up-regulated genes when comparing the rest of tNBs with cluster 7. (**D**) Temporal identity genes are not enriched in E93^+^ or Imp^+^ tNBs.

**Figure 6 - figure supplement 1:**
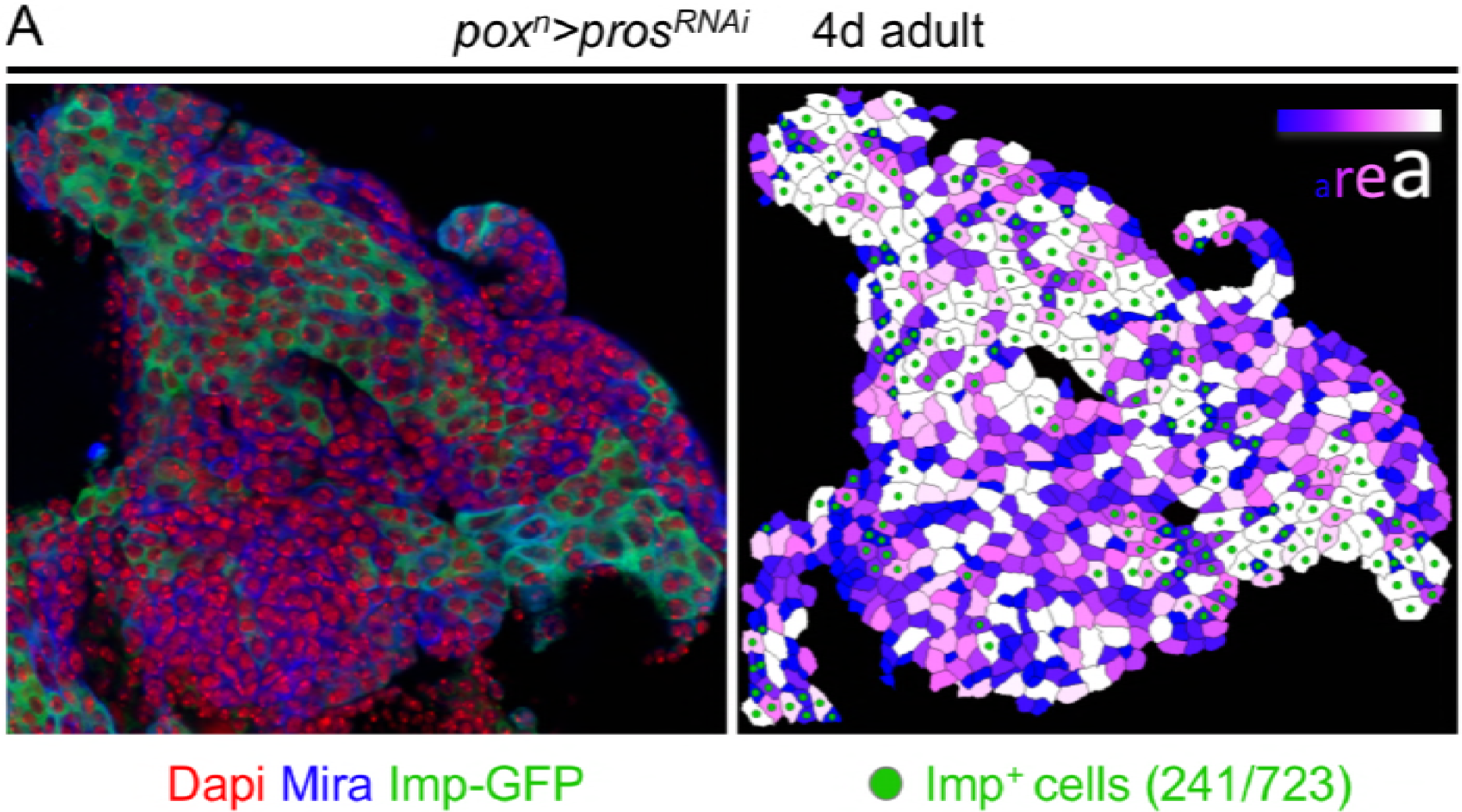
Measurement of tNB size. Segmented confocal section of a tumor using Tissue Analyzer. Segmented cells are color-coded relative to their size. Imp-GFP+ cells represent Chinmo^+^Imp^+^ tNBs. GFP^-^ tNBs represent Syp^+^ tNBs.

